# Disentangling temporal and rate codes in primate somatosensory cortex

**DOI:** 10.1101/2022.09.19.508566

**Authors:** Thierri Callier, Thomas Gitchell, Michael A. Harvey, Sliman J. Bensmaia

**Affiliations:** Committee on Computational Neuroscience, University of Chicago, Chicago, IL; Department of Organismal Biology and Anatomy, University of Chicago, Chicago, IL; Department of Medicine, University of Fribourg, 1700 Fribourg, Switzerland; Neuroscience Institute, University of Chicago, Chicago, IL

## Abstract

Millisecond-scale temporal spiking patterns encode sensory information in the periphery, but their role in cortex remains controversial. The sense of touch provides a window into temporal coding because tactile neurons often exhibit precise, repeatable, and informative temporal spiking patterns. In somatosensory cortex (S1), for example, responses to skin vibrations exhibit phase-locking that faithfully carries information about vibratory frequency. However, the respective roles of spike timing and rate in frequency coding are confounded because vibratory frequency shapes both the timing and rates of S1 responses. To disentangle the contributions of these two neural features, we measured S1 responses as animals performed a frequency discrimination task, in which differences in frequency were accompanied by behaviorally irrelevant variations in amplitude. We then assessed the degree to which the strength and timing of S1 responses could account for the animals’ performance on the task. First, we showed that animals can discriminate frequency, but their performance is biased by amplitude. Second, rate-based representations of frequency are susceptible to changes in amplitude, but in ways that are inconsistent with the animals’ behavior, calling into question a rate-based code for frequency. In contrast, timing-based representations are impervious to changes in amplitude, also inconsistent with the animals’ behavior. We account for the animals’ behavior with a model wherein frequency coding relies on a temporal code, but frequency judgments are biased by perceived magnitude. Our results constitute further evidence for the role of millisecond-scale spike timing in cortex.

**Significance statement:** While neurons in the cerebral cortex are known to produce temporally patterned responses with single-digit millisecond precision, the role of this patterning remains controversial. The alternative hypothesis is that the critical feature of neuronal responses is the rate at which spikes are emitted, and the distribution of the spikes in time only matters at much slower time scales (tens or hundreds of milliseconds). To disentangle these two putative neural codes, we trained animals to discriminate the frequency of skin vibrations while recording from somatosensory cortex. We show that only a timing-based representation of frequency can account for the performance of the animals, though overall population firing rate biases their frequency judgments.

## Introduction

Neurons throughout the nervous system convey information not only in the rate at which they emit spikes but also in the distribution of these spikes across time. The somatosensory system is a fertile model to study the role of spike timing in nervous system function given the exquisitely precise temporal patterning in the spiking responses of neurons across large swaths of its neuraxis. For example, tactile nerve fibers produce responses to vibrations and textures whose millisecond-precision timing carries information about the frequency composition of the vibration or the identity of the texture and shapes the evoked percept ^1–6^. Such informative temporal spiking patterns are also observed in the central nervous system (CNS), including the cuneate nucleus ^7^, thalamus ^8,9^, and somatosensory cortex (S1) ^3,10–18^. However, the role of spike timing in the CNS is more difficult to establish because, unlike tactile nerve fibers, which can be divided into a handful of clearly delineated neuronal classes, central neurons exhibit highly heterogeneous responses so a large population of such neurons can, in principle, encode any spatiotemporal input pattern in its rate response ^10^.

S1 responses to sinusoidal vibrations phase lock to each cycle ^11–16,18^, so vibratory frequency can be inferred from this temporal patterning, leading to the hypothesis of a temporal code for frequency in cortex ^14^, as had been proposed for the nerve ^5^. However, changes in frequency are accompanied by changes in firing rate, and this rate-based signal – under some conditions at least – is also highly informative about frequency and has been put forth as an alternative neural code ^13,16,18^. The viability of the rate code breaks down at high frequencies (> ∼80 Hz) as the relationship between firing rate and frequency breaks down ^11^. Even at the low frequencies, the rate code for frequency is, in principle at least, susceptible to large amplitude-dependent distortions given the strong and nearly universal impact amplitude has on S1 firing rates. To further muddle the issue, changes in frequency not only change the perceived pitch of a vibration – a sensory continuum analogous to its auditory counterpart –, but also its perceived intensity ^19^, likely resulting from the frequency-dependent changes in cortical rates.

The objective of the present study was to disentangle the various factors that shape pitch perception to assess the contribution of spike timing to neural coding in cortex. To this end, we trained monkeys to discriminate the frequency of sinusoidal vibrations delivered to their fingertips in the presence of concomitant variations in vibratory amplitude. The animals thus had to learn to judge pitch independently of sensory magnitude. While the animals performed this task, we measured the responses evoked in S1, including Brodmann’s areas 3b, 1, and 2. First, we found that the animals could learn to discriminate frequency, but their frequency judgments exhibited amplitude-dependent biases. Second, frequencies decoded from population response rates also exhibited amplitude biases, but these were inconsistent with those of the animals. Third, temporal patterning in S1 responses was highly informative about frequency. However, the timing-based signal was impervious to variations in amplitude so could not account for the biases in the animals’ frequency judgments. A model wherein frequency is encoded in spike timing but pitch judgments are biased by perceived magnitude can account for the animals’ behavior. We conclude that millisecond-scale spike timing contributes to sensory coding in cortex.

## Results

### Monkeys can discriminate frequency despite variations in amplitude

Human subjects can discriminate the frequency of vibrations applied to the skin even in the presence of concomitant variations in amplitude ^11,20^. Under these experimental conditions, subjects cannot exploit frequency-dependent differences in magnitude to perform the task and must rely on differences in pitch. To probe monkeys’ ability to judge frequency independently of amplitude, we had them perform a frequency discrimination task (Figure 1). In one version of the task, animals were sequentially presented with a pair of vibrations and reported which of the two was higher in frequency. In another version of the task, they were presented with a single vibration and classified it as higher or lower than a separatrix, which they learned via trial and error over a few hundred trials. Both paradigms yielded similar results. In both cases, the amplitude of each stimulus varied from trial to trial to reduce or eliminate the reliance on frequency-dependent differences in perceived magnitude to perform the task, as these were drowned out by uninformative, amplitude-dependent variations in perceived magnitude.

**Figure 1:**
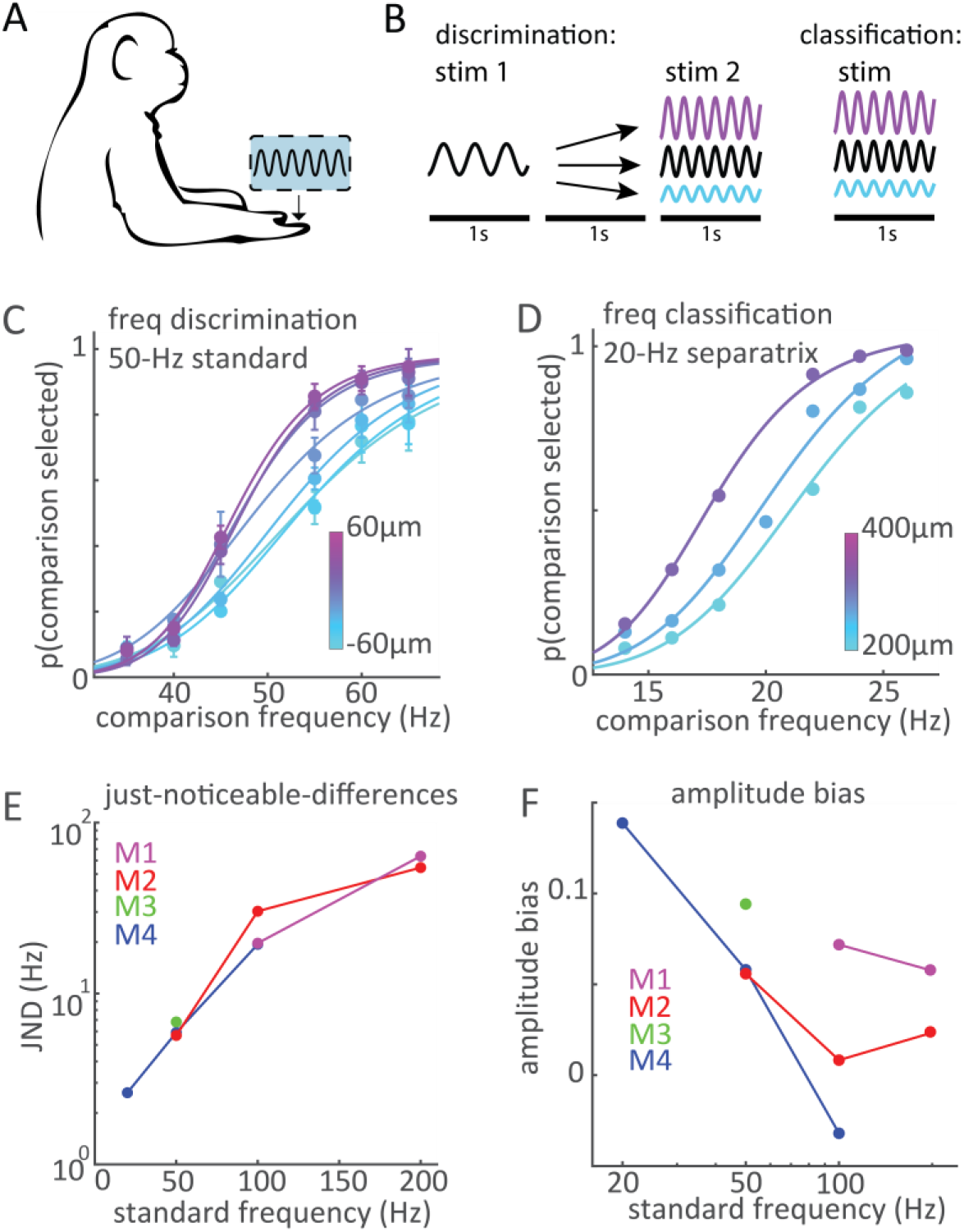
Behavioral task. A| Sinusoidal vibrations were delivered via a punctate probe driven by a shaker motor. B| In the discrimination task (left), two vibrations differing in frequency were sequentially presented on each trial and the animal judged which was higher in frequency. The amplitude of both the standard and the comparison varied from trial to trial, to preclude performing the task based on frequency-dependent differences in sensory magnitude. In the frequency classification task (right), a vibration at one of six frequencies was presented on each trial and the animal judged whether it was higher or lower than a separatrix, learned through trial and error. The amplitude of the vibration varied randomly from trial to trial to preclude performing the task based on intensity cues. C| Mean performance of monkeys M2 and M3 on the discrimination task with the 50-Hz standard. Different colors denote different amplitude differences between the comparison and standard stimulus. Psychometric functions for all standards are show in Figure S 1A. D| Performance of monkey M4 on the frequency classification task with the 20-Hz separatrix. Psychometric functions for all separatrices are show in Figure S 1B. E| Just noticeable differences (JNDs) for all animals and standards/separatrices. F| The amplitude bias – the mean pairwise difference between p(comparison selected) within the same frequency but at different amplitude conditions – gauges the tendency to judge higher-amplitude vibrations as being higher in frequency.

With both paradigms, we found that animals were able to discriminate frequency in the presence of fluctuations in amplitude (Figure 1C, D; Figure S 1A,B). Indeed, as the comparison frequency increased, the likelihood of judging it as higher in frequency increased as well. Tactile frequency perception was more acute at the low than high frequencies, with JNDs increasing systematically as the standard frequency increased (Figure 1E). JNDs tended to increase faster than did the standard frequency, leading to an increase in Weber fraction with frequency (Figure S 1C), as has been found in humans ^21^. JNDs and Weber fractions derived from the monkeys’ performance were similar to their human counterparts ^15,21–23^.

Even after extensive training, however, animals retained a bias toward judging higher-amplitude vibrations as higher in frequency, as evidenced by the spread in the psychometric functions (Figure 1F, upward shift as curves progress from cyan to purple). The amplitude bias was more pronounced for some animals than others, was stronger at some frequencies than others and for more difficult comparisons (Figure S 1D). Similar amplitude biases have also been observed in the pitch judgments of humans ^20^.

### S1 responses depend both on vibratory frequency and amplitude

Next, we investigated the dependence of neuronal responses in S1 on the frequency and amplitude of sinusoidal skin vibrations. First, we reanalyzed the responses of 211 single units in S1 to vibrations, recorded sequentially from two animals (M5, M6) using electrode drives (previously reported in ^11,24^) as well as the multi-unit S1 responses, recorded simultaneously from one animal (M4) using chronically implanted arrays (Figure 2A). The animals were not performing a behavioral task during these neurophysiological measurements, but the stimulus set tiled the space of vibratory frequencies and amplitudes more systematically than did that used in the behavioral experiments (discussed below). In this study, we draw on data obtained in several experiments with different animals, tasks, and types of neural recordings (single vs. multi-units). Table S 1 summarizes the contribution of each data set to each analysis and figure.

**Figure 2.**
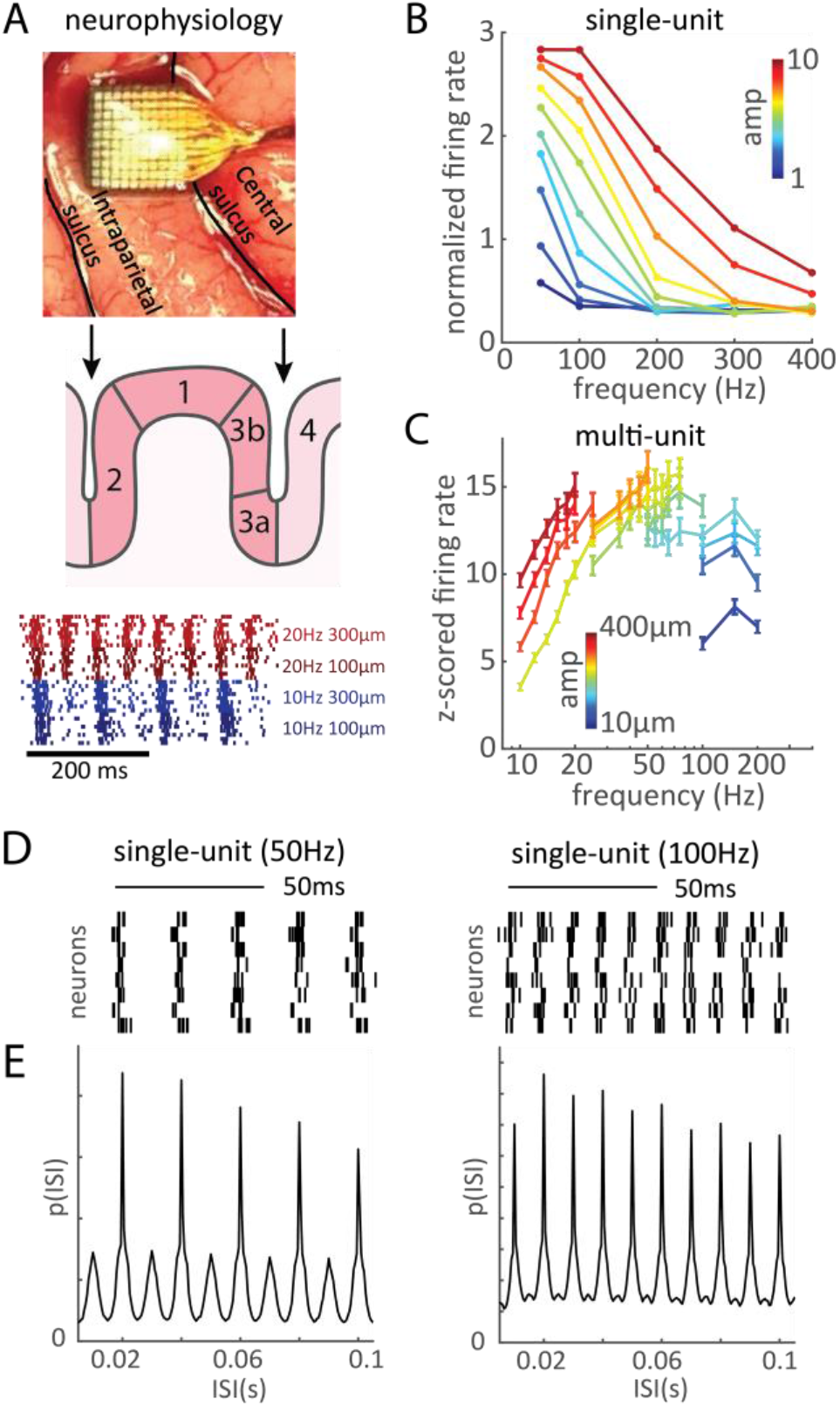
Dependence of firing rate and spike timing on frequency. A| Top: Placement of an electrode array in the hand representation of Brodmann’s area 1 of S1. Middle: Using electrode microdrives, we recorded the responses of neurons in Brodmann’s areas 3b, 1, and 2. Bottom: Responses of an S1 multi-unit from monkey M4 to vibrations at 2 frequencies and amplitudes. The response rate is higher at the higher frequency, but also at the higher amplitude. In contrast, the temporal patterning is consistent across amplitude. B| Firing rate vs. frequency for single-unit S1 responses (M5), averaged across the population. For each neuron, responses to each stimulus were normalized by the average response of the neuron across all stimuli. Different colors denote different amplitudes (ranked from lowest to highest). C| Firing rate versus frequency for a representative multiunit channel with a strong rate-frequency relationship (M4). Rate rises steeply with frequency at the low but not high frequencies. Different colors denote different stimulus amplitudes. D| Response of eight S1 neurons to a 50-Hz (left) and 100-Hz (right) vibration. E| Inter-spike interval distribution of the population response to vibrations at 50 and 100 Hz.

Examination of the rate-intensity function revealed that, while almost all S1 neurons responded vigorously to vibrations applied to their receptive fields (Figure 2A-C), the dependence of the response on frequency varied across the population. Indeed, some S1 neurons exhibited a strong sensitivity to frequency (Figure 2C) whereas others did not (Figure S 2A-C). For frequency-sensitive neurons, frequency dependence tended to be more prominent at the low than high frequencies (Figure 2B,C). Over the range of frequencies tested, no S1 responses were tuned to frequency, peaking at one frequency and trailing off in both directions, unlike what has been found in mouse S1 ^25^. The responses of all neurons increased with amplitude at all frequencies (Figure S 2D-F).

S1 responses also phase locked with the vibration, producing one or more spikes within a restricted phase of each stimulus cycle (Figure 2D). This phase-locking produced characteristic distributions of inter-spike intervals (ISIs), which were most distinctive at low frequencies and became less so at higher frequencies (Figure 2E, Figure S 3). As expected, phase-locking tended to be stronger in single-unit than multi-unit responses, likely because the latter are liable to reflect phase-locked responses of multiple neurons at different phases or responses that are phase-locked combined with responses that are not.

### Decoded frequency is biased by amplitude for rate-based but not timing-based decoders

Having established that both the rate and timing of S1 responses are dependent on vibratory frequency, we assessed the degree to which the frequency of a vibration could be inferred from these two features of the population response.

To decode vibratory frequency from rates, we projected the firing rates of the neural population onto a lower dimensional subspace using principal components analysis (PCA) then performed a linear discriminant analysis (LDA) on the responses projected onto this reduced subspace. We found that amplitude variation introduced systematic errors into rate-based decoding, with a bias toward selecting the lower-amplitude vibration as higher in frequency (Figure 3A). Note that the amplitude bias from these rate-based decoders is opposite in sign to that observed in the monkeys’ behavior. The performance of rate-based decoders did not improve with larger sample sizes (Figure 3B), nor did the amplitude bias decrease (Figure 3C). In this analysis, we decoded frequency from steady-state responses, excluding the large burst of activity at stimulus onset (Figure S 4A). As different epochs of the rate response have been found to be differentially informative about frequency ^13^, we assessed whether the response epoch starting from onset might yield more reliable frequency decoding performance and found that it did not (Figure S 7A).

**Figure 3.**
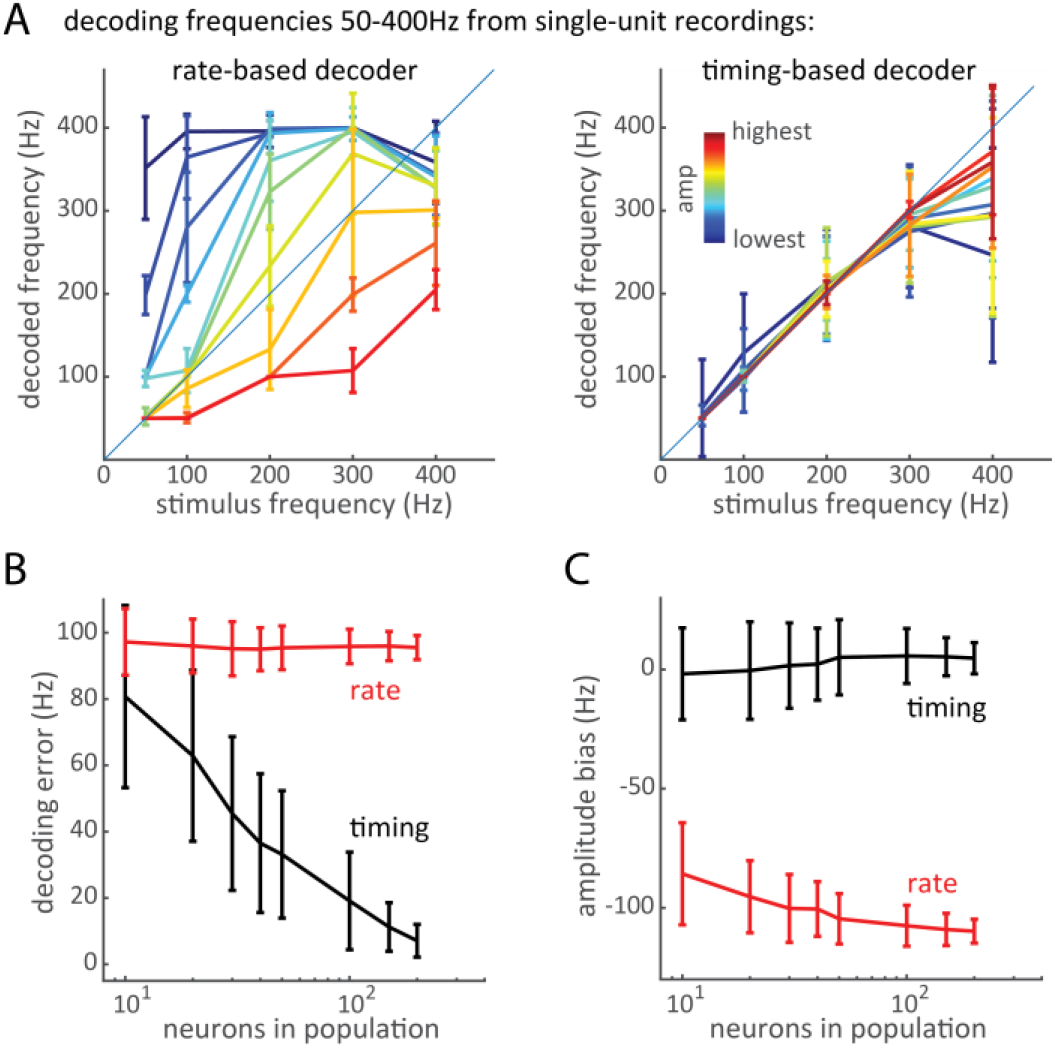
Decoding frequency from the rates and temporal patterning in the responses of single units. A| Decoded vs. actual frequency for the rate decoder and the timing-based decoder. At each frequency, ten amplitudes were used, each denoted by a different color (corresponding to its rank). Amplitude variations introduce systematic errors when decoding frequency from rate but not timing. B| Mean absolute decoding error as a function of the number of neurons in the synthetic population of responding neurons. The timing decoder improves as the population size increases. C| Amplitude bias, a measure of the difference in decoded frequency due to amplitude (i.e. of the spread between amplitude curves, see methods), as a function of population size. In panels A-C, population responses were constructed by random sampling, and error bars represent the standard deviation across 200 iterations. Panel A shows results with a population of 100 neurons.

To decode frequency from timing, we computed a template ISI distribution for each frequency (using the responses of a population of phase-locked neurons across all amplitudes, see methods) then found the best match (yielding the highest correlation) for ISI distributions derived from the responses on left-out trials. Note that the timing decoders were designed to gauge the informativeness of spike timing independent of spike rate. We found that, in contrast to their rate-based counterparts, timing-based decoders achieved precise classification with little to no bias (Figure 3A-C). The performance of the timing-based decoder was somewhat poorer at high frequencies, mirroring behavioral performance (Figure 1E, Figure S 1) and reflecting the weaker timing signal as frequency increases (Figure S 3). When the timing of the spikes was shuffled (thereby preserving rates but destroying phase-locking), the timing-based decoder failed, confirming its reliance on spike timing (Figure S 8).

Analysis of the single unit responses to vibrations spanning a wide range of frequencies reveals that the frequency signal in the firing rates is highly contaminated by variations in amplitude whereas that in the phase-locking is not. The advantage of this stimulus set is that neurons were probed with vibrations that spanned a wide range of frequencies. The disadvantage is that the range of amplitudes differed across frequencies because the highest achievable amplitude at 400 Hz yields a barely perceptible vibration at 50 Hz. To verify that our results were not an artifact of different amplitude ranges, we analyzed multi-unit S1 responses to vibrations whose amplitudes were identical across frequencies but, by necessity, spanned narrower ranges of frequency. We first focused on vibrations in the flutter range (10-20 Hz) to replicate previous findings that rate-based signals are highly informative over this range ^13,17,18^. To match these previous studies to the extent possible, we initially restricted the analysis to trials on which the amplitude of the two stimuli in a pair was fixed, thereby eliminating any confounding effects of amplitude variations on the frequency signal carried by the firing rates. For these fixed-amplitude trials, the decoded frequency systematically matched the actual frequency (Figure 4A left panel), consistent with previous results (cf. ^13,17,18^). When variations in amplitude were introduced, however, rate-based decoders exhibited a strong amplitude bias (Figure 4A middle panel, Figure S 9).

**Figure 4.**
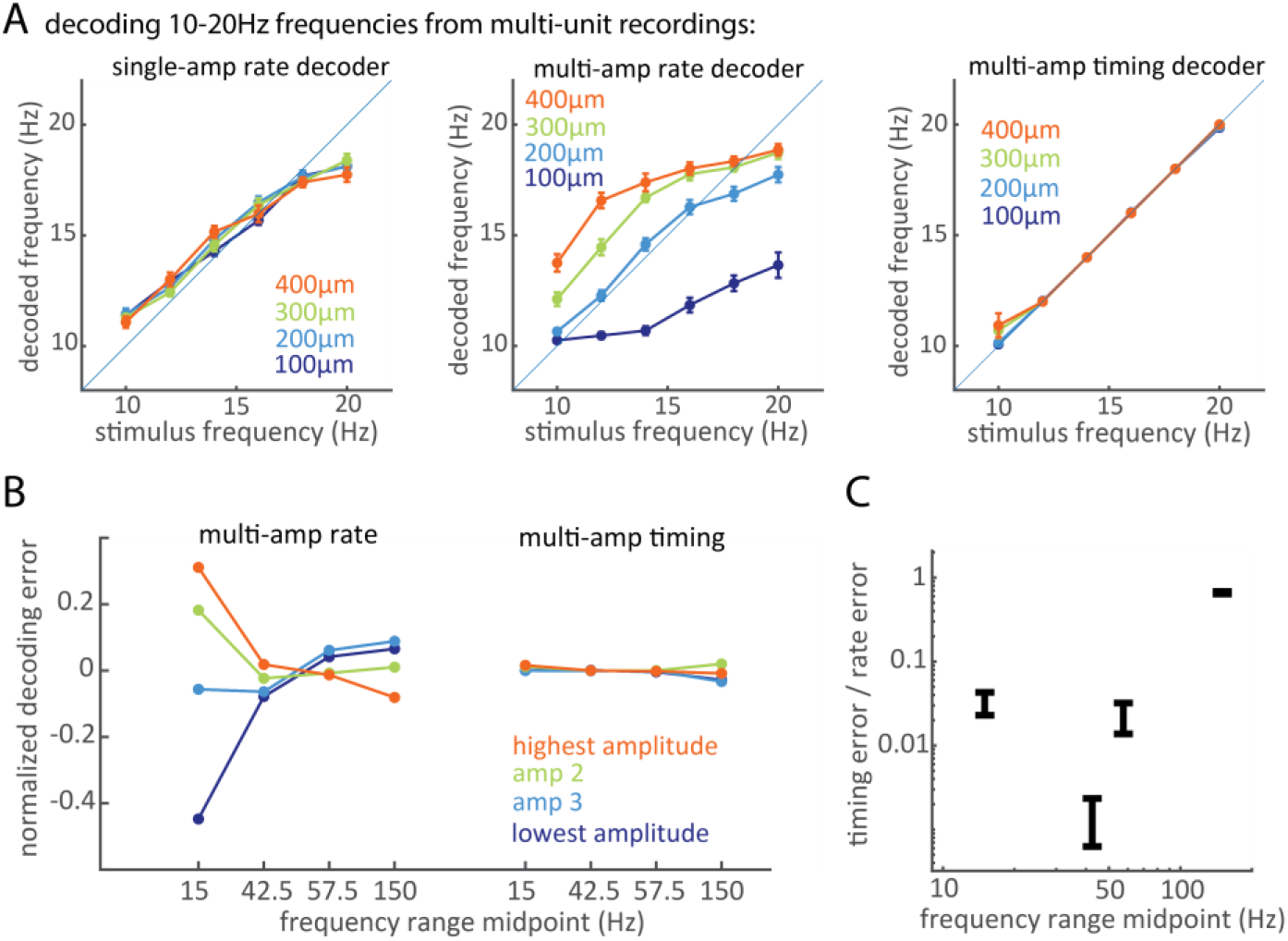
Decoding frequency from the rates and temporal patterning in the responses of multi-units. A| Decoded vs. actual frequency in the low frequency range, using (left) a rate-based decoder with constant amplitudes (a rate decoder was trained and tested at each amplitude separately), (middle) a rate-based decoder with all amplitudes (a single rate decoder was trained and tested with all amplitudes), and (right) a timing decoder with all amplitudes. See Figure S 9 for decoding in other frequency ranges. B| Decoding error broken down by amplitude for the rate decoder (left) and the timing decoder (right). The sign of the error (decoded frequency – actual frequency) is preserved: positive errors denote systematic overshoots and vice versa. Errors from each frequency range were computed separately for each amplitude, averaged across frequencies in that range, and normalized by the frequency range. C| Ratio of the decoding error of the timing-based decoder to that of the rate-based decoder. Timing decoder outperforms rate decoder at all frequencies. Unlike in panel B, only the magnitudes of the errors are compared without considering the direction of error. The absolute values of decoding errors were averaged across all frequencies and amplitudes in each frequency range.

The amplitude bias was observed for all frequency ranges (10-20 Hz, 35 – 50 Hz, 50-65 Hz, and 100-200 Hz), but its direction differed in different frequency ranges (Figure 4B). At the low frequencies, rate-based decoders tended to underestimate the frequency of low-amplitude stimuli and overestimate that of high-amplitude stimuli, but the reverse was true at the high frequencies. Thus even when the amplitude range was consistent across frequencies, neuronal firing rates yielded poor estimates of frequency in the presence of variations in amplitude.

In contrast, timing-based decoders yielded near perfect performance over the flutter range (Figure 4A right panel) and outperformed their rate-based counterparts (with errors one or more orders of magnitude smaller) across all but the highest frequencies, where the performance of the two decoders was comparable (Figure 4C). The poor performance of timing-based decoders at the high frequencies reflects the degradation of timing in the multi-unit response at those frequencies (Figure S 3). The timing-based decoders were also far less susceptible to variations in amplitude than were rate-based decoders across the frequency range (Figure 4B). In other words, the phase-locking in the response yields accurate estimates of frequency despite the variations in amplitude. The timing-based decoders outperformed the rate-based ones even though the rate-based decoding involved parameter optimization (LDA weights) and timing-based decoding did not entail any parameter fitting.

### Rate-based and timing-based decoders cannot account for behavioral performance

The above classification analyses gauge the tendency of rate-based and timing-based decoders to reconstruct vibratory frequency across repeated stimulus presentations but their outcome cannot be directly compared to the psychophysical performance of the animals. To fill this gap, we analyzed S1 responses recorded while the animals performed the discrimination or classification task and simulated the tasks with rate-based and timing-based decoders. In brief, each decoder was trained on neuronal responses to all but one stimulus then tested on the held-out responses. To simulate the two-alternative forced-choice task, frequency was decoded from the single-trial response in each interval and the higher frequency was selected. We then computed the proportion of times the decoded frequency of the comparison stimulus was higher than that of the standard for each frequency/amplitude combination (Figure 5A, Figure S 10). In the classification task, we computed the proportion of times of times the decoded frequency exceeded the separatrix (Figure S 10). We also implemented a decoder that simply selected the stimulus that evoked the higher rate as being higher in frequency (“greater-rate” decoder, Figure S 11A, B).

**Figure 5.**
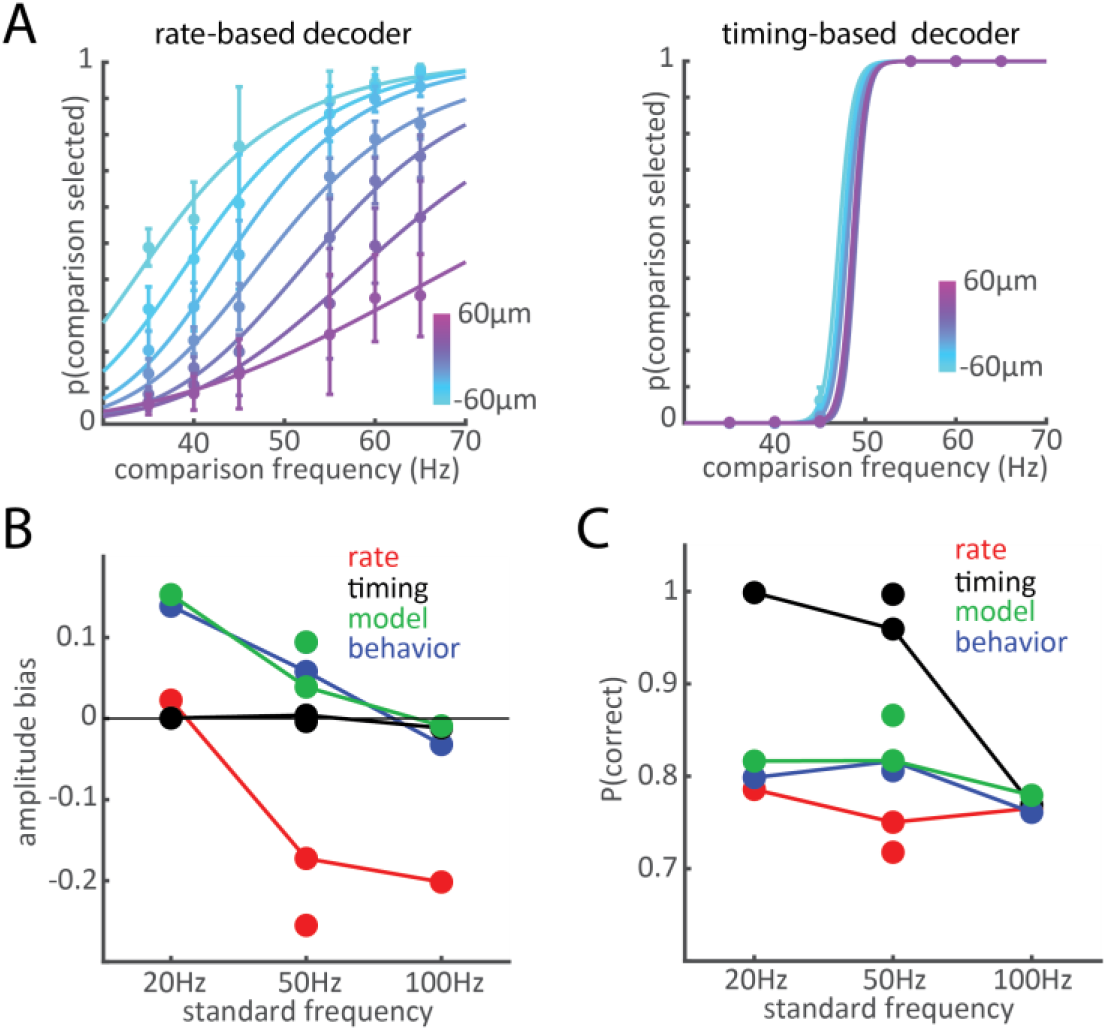
Simulation of the behavioral tasks with rate-based and timing-based decoders. A| Discrimination performance of a rate decoder (left) or timing decoder (right) based on the neural responses of M3 recorded during frequency discrimination with a 50-Hz standard. Different colors denote different amplitude differences between comparison and standard. The neurometric functions derived over other frequency ranges for the frequency discrimination and classification tasks are shown in Figure S 10. B| Amplitude biases of the rate decoder, timing decoder, and rate-biased timing model, along with the animals’ behavioral bias. A positive bias indicates a tendency to select high-amplitude stimuli as higher in frequency. The timing decoders exhibits a negligible amplitude bias in all cases. The rate decoders exhibit substantial biases that differ from those of the animals in magnitude and sign. The rate-biased timing model exhibits comparable biases to those of the animals. C| Average discrimination/classification performance for the decoders and animals. The animals slightly outperformed the rate decoders, but the timing decoders outperformed the animals. The performance of the model – which posits that frequency judgments are based on timing but biased by rate – approximates that of the animals.

Consistent with the classification analyses (Figure 4), the rate-based decoders exhibited strong amplitude biases whereas the timing decoders did not (Figure 5A, B, Figure S 10). The overall performance of the rate-based decoders could be improved by including more neural factors (PCs, Figure S 12) but the amplitude bias persisted. Furthermore, rate-based decoders exhibited a tendency to judge the lower-amplitude vibration as higher in frequency (Figure 5A, B, Figure S 10) whereas the monkeys maintained a bias toward higher amplitudes in almost every case (the only exception being M4 with the 100-Hz separatrix; Figure 5B, Figure S 1A, B). The biases of the rate-based decoders are thus of the opposite sign as those of the animals.

The timing-based decoders yielded far better performance than the animals at the low frequencies (Figure 5C), consistent with previous findings ^18^. They also were far less susceptible to variations in amplitude than were monkeys (Figure 5B), consistent with the classification analyses (Figure 4). In summary, rate-based decoders made erroneous predictions about the monkeys’ biases and timing-based decoders were more accurate and less biased than the animals. We repeated these analyses with neuronal populations broken down by cortical field and found similar results across fields (see Supplemental Results).

### A temporal code with a rate bias can account for behavioral performance

If the animals based their judgments solely on differences in magnitude, they would select the stimulus that evokes the strongest response across the population, as sensory magnitude is determined, at a first approximation, by population rate ^11,26^. The magnitude discrimination strategy yields neurometric functions that feature a positive amplitude bias, in the same direction as its behavioral counterpart but much larger in magnitude (Figure S 11A, B). The animals’ frequency judgments are thus intermediate between the unbiased judgments predicted from timing-based decoders and the strongly biased judgments predicted from the magnitude discrimination strategy. With this in mind, we explored the possibility that frequency judgments rely primarily on temporal codes and that amplitude biases reflect a tendency of the animals to judge vibrations that feel more intense as being higher in frequency. To test this hypothesis, we implemented a model whose output was determined by a weighted combination of the output of a timing-based decoder and that of the “greater rate” decoder. We found that the predictions of this rate-biased timing model were consistent with behavior both in terms of overall performance (Figure 5C) and bias (Figure 5B, Figure S 11C, D).

In conclusion, rate-based decoders exhibit an amplitude bias that is not consistent with its behavioral counterpart in the frequency discrimination task. In contrast, timing-based decoders exhibit no amplitude bias, also inconsistent with behavior. However, a hybrid model that posits that the animals rely on temporal spiking patterns for pitch perception but are biased by sensation magnitude in their pitch judgments accounts for performance.

## Discussion

We showed that animals can judge the pitch of tactile vibrations in the presence of concomitant and uninformative variations in sensation magnitude arising from variations in stimulus amplitude. However. the animals maintained an amplitude-dependent bias in their frequency judgments despite extensive training. Seeking the neural basis for vibrotactile pitch discrimination, we found that frequencies decoded from neuronal firing rates depended on vibratory amplitude in fundamentally different ways than did the corresponding pitch judgments. In contrast, decoding frequency from the temporal patterning in the response – reflecting phase-locking to the vibration – yielded more reliable and nearly amplitude-independent pitch discrimination judgments. We can account for the monkeys’ behavior with a model wherein pitch perception relies on a temporal code in S1 but pitch discrimination judgments are biased by sensation magnitude, for which population firing rate is the neural correlate.

### Vibratory frequency and amplitude shape touch sensations in overlapping ways

A major challenge in establishing a neural code is that stimulus dimensions are typically not encoded independently. In the sense of touch, for example, the neural representations of texture and scanning speed are tangled in the somatosensory nerves ^6,27^. Texture and speed signals are untangled via downstream processing ^28,29^, a process that culminates in a stable representation for texture but not speed ^27,30,31^. The two parameters that define a sinusoidal vibration – frequency and amplitude – modulate the resulting experience in overlapping ways. While vibratory frequency is the major determinant of pitch and vibratory amplitude is the main determinant of perceived magnitude, magnitude is dependent on frequency ^19^ and pitch is dependent on amplitude ^20,32^. Frequency discrimination judgments can thus, in principle, reflect changes in both pitch and magnitude. One approach to disentangle these two perceptual correlates of vibratory frequency has consisted in equating perceived magnitude in preliminary experiments, then having subjects discriminate the frequency of intensity-matched vibrations. However, the matching is only approximate and so is not guaranteed to eliminate intensity cues. The confounded effects of the two stimulus parameters on the evoked percepts complicates inquiries into the neural codes that mediate both pitch and magnitude perception ^33,34^.

### The temporal patterning in and strength of the S1 response drive frequency discrimination performance

We show that temporal spiking patterns carry robust information about vibratory frequency whereas rate responses carry more ambiguous frequency signals. However, timing-based decoders are inconsistent with the animals’ behavior in that their frequency discrimination performance is more acute and less susceptible to amplitude biases. We propose that the rate of S1 neurons does not play a role in pitch perception *per se*. Rather, animals exhibit a bias to judge more intense vibrations as being higher in frequency. The rate-biased timing model accounts for the animals’ behavior but can also account for the amplitude biases observed in the pitch judgments of human subjects (cf. ref. ^20^). Over the range of frequencies tested here, higher frequencies give rise to more intense percepts, so more intense percepts are liable to be interpreted as stemming from higher frequencies. At vibratory frequencies above around 300 Hz, increases in frequency lead to a decrement in perceived magnitude ^19^, due to a decreased sensitivity of tactile nerve fibers, in particular Pacinian Corpuscle associated fibers ^35^. Over this range, humans exhibit a reverse amplitude bias, with higher amplitudes interpreted as lower in frequency ^20^. This observation is consistent with the rate-biased timing model if one assumes that the sign of the amplitude bias is determined by the sign of the impact of frequency on sensory magnitude.

According to the rate-biased timing model, then, pitch is encoded in the timing and the rates contaminate pitch judgements at a cognitive rather than sensory level: when uncertain, the animal selects the more intense stimulus. An alternative interpretation of the results is that the downstream neural circuits that extract pitch from spike timing are less than ideal. Indeed, timing-based decoders are implemented using Fourier analysis ^18^ or ISI distribution template matching (here) and thus do not take into consideration how stimulus information is extracted from temporal spiking patterns. Neural circuits are likely to be less reliable than the computational approaches used here to estimate timing, leading to poorer frequency discrimination. In fact, these circuits may also yield frequency estimates that are contaminated by rate (see below).

Both the rate-base code and the rate-biased timing model predict that rates would co-vary on a trial-to-trial basis with the animals’ choice. For the rate-biased timing model, a vibration that evoked a higher firing rate would tend to be perceived as higher in frequency. While this relationship has been previously observed ^18^, we found it to be inconsistent across animals: One animal exhibited the expected relationship, the other did not (see Supplementary Results). This inconsistency casts doubt on the interpretability of these analyses in the present context.

### Temporal coding shapes the perception of non-periodic stimuli

Vibratory frequency discrimination is a fruitful window into the role of spike timing in perception given the simplicity of the stimulus space and the relative ease with which animals can be trained to perform the task. The periodicity of the vibrations further simplifies the description of the stimulus and the underlying temporal code. However, the principles revealed through the lens of frequency discrimination extend to stimuli of arbitrary complexity. For example, monkeys’ ability to discriminate the frequency of non-periodic vibrations can also be accounted for using a temporal code ^12,13^. Each stimulus event is encoded in a spike or burst distributed over a population of cortical neurons, and the number of spikes/bursts over the stimulus interval forms the basis for the animals’ judgments. The timing-based decoder used here – ISI-template matching – could not be applied to non-periodic stimuli, but analogous decoders – based for example on the median ISI – may. The temporal coding hypothesis further predicts that different aperiodic vibrations at the same nominal frequency are liable to be discriminable from one other. Consistent with this hypothesis, humans can distinguish diharmonic vibrations that differ in the phase of one component relative to the other ^36^, suggesting that the distribution of stimulus peaks over time, not only the rate at which they occur, shapes the evoked percept.

Similarly, textures scanned across the skin elicit texture-specific vibrations in the skin ^37–39^, which in turn elicit texture-specific spiking patterns in tactile nerve fibers^2,6^ and cortical neurons^3,10^. These millisecond-scale temporal spiking patterns, often complex and non-periodic, convey rich information about texture and seem to play an important role in shaping the textural percept. Converging evidence using simple sinusoids – whose parameters are under experimental control – and textures – which offer the advantage of being ethological and arbitrarily complex – supports the conclusion that spike timing in somatosensory cortex plays a key role in tactile coding.

### Central neurons decipher temporal codes by differentiating their inputs

If spike timing at millisecond time scales carries stimulus information, how is this information extracted by downstream neurons? As mentioned above, approaches to infer stimulus properties from temporal spiking patterns do not take into account biological implementation. As the body of evidence for temporal coding of frequency grows ^1,3–6,11^, so does the need to explain how these timing-based signals can be integrated with rate-based ones. Indeed, at the highest level of somatosensory processing (secondary somatosensory cortex, S2), temporal patterning has all but disappeared ^12^, implying that information carried in spike timing has been converted into a rate-based representation.

We propose that this timing-to-rate conversion is mediated by neural computations that confer to individual neurons a preference for specific temporal features in their inputs. Indeed, reverse correlation reveals that neurons act as temporal filters on their afferent input ^24^. Furthermore, the filters comprise excitatory and inhibitory lobes, essentially implementing a temporal differentiation, which confers to the neurons a preference for time-varying inputs over constant inputs. The width and relative strength of the excitatory and inhibitory lobes vary idiosyncratically from neuron to neuron, which confers to different neurons preferences for different temporal features in their inputs. This temporal filtering, prominent in S1 ^24^, begins to emerge at the first stage of tactile processing along the medial lemniscus-dorsal column pathway, namely the cuneate nucleus ^7^. This gradual process of converting temporal patterns into rate-based signals is not complete in S1, as evidenced by the poor rate-based representations of frequency. Consistent with this hypothesis, the informativeness of temporal spiking patterns decreases even within somatosensory cortex: as one progresses from area 3b, to area 1, to area 2, the incidence of precisely timed responses decreases ^3,11^.

Note that, if somatosensory neurons acted as sinusoidal filters whose period varies from neuron to neuron, a population of such neurons would effectively implement a Fourier-like spectral decomposition of skin vibrations. Because these temporal filters are not sinusoidal (but rather take on highly idiosyncratic shapes ^7,24^), their output does not provide such an unbiased spectral decomposition. This offers an alternative to the rate-biasing proposed here: The process of extracting pitch from timing may introduce a rate-bias, reflected as an amplitude bias in the evoked pitch percept. For example, the extent that the excitatory and inhibitory lobes are imbalanced, the output of the filter will tend to covary with input rate ^24^. Regardless, somatosensory neurons in the central nervous system exhibit properties which, in principle, could extract stimulus information from temporal spiking patterns.

## Conclusions

S1 firing rates cannot straightforwardly account for the frequency discrimination judgments of monkeys and humans. A rate-based code for frequency makes erroneous predictions about how variations in amplitude influence frequency discrimination judgments. While temporal patterning in S1 can account for the ability to discriminate pitch, a temporal code for pitch makes the erroneous prediction that pitch judgments are impervious to differences in amplitude. We propose that pitch is encoded in temporal spiking patterns but that pitch judgments are biased by sensory magnitude.

## Methods

We report on data obtained in two separate studies. The first study yielded acute recordings from somatosensory cortex using a multielectrode Microdrive (previously reported in ^11,24^). The second yielded chronic recordings using implanted electrode arrays (never previously published). All procedures were approved by the University of Chicago Animal Care and Use Committee.

### Neurophysiological recordings

#### Acute recordings

We obtained extracellular recordings from the postcentral gyri in four hemispheres of two macaque monkeys (M5 and M6), using previously described techniques ^11^. In brief, two to four resin-coated electrodes (FHC Inc.) were inserted into the postcentral gyrus using a microdrive (NAN Instruments) normal to the cortical surface. Brodmann’s areas 3b, 1, and 2 were identified based on stereotypical RF properties and relative RF locations as one progresses along the gyrus and the posterior bank of the central sulcus. We recorded from neurons whose RFs were located on the distal pads of digits 2–5. On every second day of recording, the electrode array was shifted 200 µm along the postcentral gyrus until the entire representation of digits 2–5 had been covered. Recordings were obtained from S1 neurons that met the following criteria: (1) action potentials were well isolated from the background noise, (2) the RF of the neuron was on the glabrous skin, and (3) the neuron was clearly driven by light cutaneous stimulation.

#### Chronic recordings

Four male Rhesus macaques (*Macaca mulatta*), 7-13 years old and weighting 9-12kg, participated in the behavioral study. M1 and M2 were implanted long (4+ years) before the beginning of the study, precluding any neural data collection due to degraded signal quality. Only behavioral data are reported for these two animals. M3 and M4 were implanted shortly after they learned to perform the behavioral tasks, so both neurophysiological and behavioral data were obtained from them. Each animal was implanted with one Utah electrode array (UEA)(Blackrock NeuroTech Salt Lake City, UT) in the hand representation of Brodmann’s area 1 in somatosensory cortex (S1) (Figure 2A). Each UEA consists of 96 1.5mm-long electrodes with tips coated in iridium oxide, spaced 400 µm apart, and spanning 4 mm x 4 mm of the cortical surface. The hand representation in area 1 was targeted based on anatomical landmarks. Given that the arrays were contiguous to the central sulcus and area 1 spans approximately 3-5mm of cortical surface from the sulcus ^40^, few if any electrodes were located in Brodmann’s area 2. Given the length of the electrodes, their tips likely terminated in the infragranular layers of the somatosensory cortex if embedded to their base, as we have previously shown in postmortem histological analysis with other animals instrumented with identical arrays ^41^.

### Tactile stimuli

#### Acute recordings

Experimental procedures have been previously described ^11^. Stimuli consisted of 1-s long sinusoidal vibrations delivered to the distal pads of the digits using a 2-mm-diameter probe, pre-indented into the skin by 0.5 mm and driven by a shaker motor (LW-132-7 Electrodynamic Shaker System, Labworks Inc.). The shaker was calibrated before each session to ensure that the stimuli were accurate and consistent across sessions. The accuracy of the stimulus traces was verified with an accelerometer (8702B50M1, Kistler Instruments Corp.) and a Laser Doppler vibrometer (LDV, Polytec OFV-3001 with OFV 311 sensor head, Polytec, Inc.). The experimental decision to pre-indent was intended to mitigate the effects of small changes in the animal’s hand position, and predicated on the fact that (1) afferent responses to pre-indentation decay within 10-20s ^42^, as does the resulting sensation, and (2) afferent responses to indentation to the skin beyond the pre-indentation are identical to those with no pre-indentation ^43^. The frequency of the vibrations was 50, 100, 200, 300, or 400 Hz, and their amplitude range depended on frequency and was constrained by the capabilities of the shaker: 34–680 µm (zero to peak) at 50 Hz, 15– 300 µm at 100 Hz, 7–133 µm at 200 Hz, 4–82 µm at 300 Hz, and 3–59 µm at 400 Hz. At each frequency, amplitudes were incremented in 10 equal logarithmic steps over the range. Vibrations were presented 5 times each in pseudo-random order.

#### Behavioral tasks/chronic recordings

Stimuli consisted of 1-sec long sinusoidal vibrations delivered at precisely controlled frequencies and amplitudes using a custom built shaker motor ^44^ driving a 1-mm-diameter probe, pre-indented 0.5 mm into the skin. The motor was regularly calibrated with an accelerometer and the LDV. Vibrations were delivered to skin locations whose somatotopic representation in area 1 was circumscribed within the array. Stimuli were delivered to the thumb pad of M3 and the index fingertip pad of M4.

##### Frequency discrimination

The animals were seated at the experimental table facing a monitor, which signaled the trial progression (Figure 1). The animals initiated a trial by directing their gaze to a fixation cross in the center of the monitor. Two vibrotactile stimuli were then sequentially delivered to the skin, each 1-sec long and separated by a 1-sec interstimulus interval. The order of the stimuli (standard, comparison) varied randomly from trial to trial. The animals’ judged which of the two stimuli was higher in frequency, indicating their response by making a saccadic eye movement to one of two response targets that appeared after the offset of the second stimulus. A trial was aborted if the animal failed to maintain fixation on the cross until the appearance of the response targets. Correct responses were rewarded with juice. Three standard frequencies were used: 50 Hz (with comparisons ranging from 35 to 65 Hz), 100 Hz (with comparisons ranging from 50 to 300 Hz), and 200 Hz (with comparisons ranging from 50 to 400 Hz). On each trial, the amplitudes of the standard and comparison stimuli varied pseudo-randomly across a wide range, to preclude performing the task based on differences in intensity instead of pitch. Amplitudes ranged from 55 and 115 µm for the 50-Hz standard and between 25 and 55 µm for the 100- and 200-Hz standards. Standards differing in amplitude were parametrically combined with comparisons differing in both amplitude and frequency and randomly interleaved in each experimental block. M1 performed the task with the 100- and 200-Hz standards, M2 performed the task with all three standards, and M3 performed it with just the 50-Hz standard. Animals performed the task with one standard until sufficient data were collected before moving on to another standard.

##### Frequency classification

The behavioral apparatus was identical to that in the frequency discrimination experiment. On each trial, a 1-sec long vibration was delivered to the skin and the animal judged whether the frequency of the vibration was higher or lower than a separatrix, learned over the first several hundred trials by trial and error (with a constant separatrix). The animal responded by making a saccadic eye movement to one of two targets. Neurophysiological data were collected when the animal reached asymptotic performance. Three separatrices were used: 20 Hz (with comparisons ranging from 14 to 26 Hz), 50 Hz (with comparisons ranging from 35 to 65 Hz), and 100 Hz (with comparisons ranging from 70 to 130 Hz). As in the discrimination task, the amplitudes of the vibrations varied widely and randomly to prevent the animal from relying on intensive cues to make frequency judgments. Amplitudes ranged from 200 to 400 µm for the 20-Hz separatrix, from 67 to 150 µm for the 50-Hz separatrix, and from 30 to 70 µm for the 100-Hz separatrix. This task was performed by M4. Each separatrix was used until sufficient data had been collected before moving on to another.

### Analysis

#### Psychophysics

##### Just noticeable differences and Weber fractions

We fit a cumulative normal density function to performance – proportion of times the comparison was selected as higher in frequency than the standard or the separatrix – vs. comparison frequency. We fit separate functions for each amplitude condition, i.e. for each combination of comparison and standard amplitudes for the discrimination task (Figure 1C, Figure S 1A) or comparison amplitude for the classification task given the absence of standard (Figure 1D, Figure S 1B). Just noticeable differences (JNDs) and Weber fractions were calculated using only trials in which both stimuli in the pair were of equal amplitude (for discrimination) or only trials in which the single comparison stimulus was at the middle amplitude (for classification), using a criterion performance of 75% correct.

##### Amplitude bias

The left- and right-ward shifts of psychometric curves for different amplitude conditions reflect the dependence of the animals’ frequency judgments on vibratory amplitude. A leftward shift indicates a bias toward judging higher-amplitude vibrations as being higher in frequency and vice versa. To quantify this bias, we computed the difference in the probability of being selected as higher in frequency (y axis values in Figure 1C, D) between every pair of comparisons with the same frequency but different amplitude conditions (e.g. the proportion of times a comparison of 55Hz at +60 µm was selected minus the proportion of times a comparison of 55Hz at +40 microns was selected, in Figure 1C). A positive value denotes a bias toward higher amplitudes, a negative value a bias toward lower amplitudes, a value of zero denotes no bias.

#### Neurophysiology

Neural responses to vibrations begin with an onset burst lasting 100-200 ms followed by a steady state response (Figure S 4A). In most decoding analyses, we analyzed the time interval from 200 to 900 ms after stimulus onset, which excludes the onset burst and encompasses most of the steady state response.

##### Standardizing neural responses in array recordings

Multiunit activity varies widely from electrode to electrode, both at baseline when no stimulus is applied, and in response to a stimulus, as might be expected given that different electrodes acquire signals from neuron groups that vary in size, sensitivity, and spontaneous activity. To compare rate responses across electrodes, we standardized stimulus-evoked responses according to the pre-stimulus baseline activity by subtracting the mean baseline rate and dividing by the standard deviation of the baseline rate across trials. That is, responses were converted to z-scores with respect to the baseline activity (Figure S 4A, B).

##### Selecting informative single or multi units

To identify single- or multi-units whose firing rate was informative about frequency, we computed the mutual information between vibratory frequency and firing rate (Figure S 5). The most informative channel from the electrode arrays were prioritized in the rate-based decoding analyses. To identify units whose timing was informative, we quantified the strength of phase locking in each spike train using vector strength (VS) (at every frequency between 10 and 400Hz) and computed the mutual information between actual vibratory frequency and the frequency that yielded the highest VS in the neuronal response. The most informative channels from the electrode arrays (M3 and M4) were selected for the timing-based decoding of multi-unit activity, and the most informative neurons were used to construct ISI templates for single-unit (M5 and M6) timing-based decoding. For both rate and timing decoders applied to the array recordings, the precise number of channels included did not substantially impact results (Figure S 6). The array results plotted throughout the paper are for decoding based on responses from the 10 most informative channels.

##### Rate-based decoding

For each unique stimulus (frequency-amplitude combination) in turn, single-unit responses were split into a test set (one response from every neuron in the population to that stimulus) and training set (responses to all other stimuli). We then performed dimensionality reduction with principal component analysis (PCA) on the training set, projected the training set onto a lower dimensional space (defined by the first N PCs, see Figure S 12), and used the resulting projections as input to a linear discriminant analysis (LDA). We then tested the classifier on the test set, also projected onto the same lower dimensional subspace. To assess the dependence of decoder performance on sample size, this procedure was repeated with different numbers of neurons to construct the population response (Figure 3B, C). We iterated the analysis 200 times for each set of conditions. To create each population of neurons of size N on each iteration, we randomly drew *N* neurons from the pool of 211 single units. A similar procedure was followed for the array recordings except that 1) responses from one trial were used as the test set and the responses to all other trials in the same experimental block were used as the training set, and 2) responses from the most informative channels were used instead of randomly drawing from the 96 channels on each array.

##### Timing based decoding

First, we pooled single-unit responses from the 50 neurons with the most informative timing (as described above) into a synthetic population response. We computed the inter-spike intervals (ISIs) in each spike train and pooled the ISIs within frequency, to obtain ISI distribution “templates.” Then, for each stimulus (frequency/amplitude combination) in turn, we generated synthetic population responses by randomly drawing *N* neurons from the population of 211 single units and randomly selecting one spike train from each of those neurons. We then computed the ISI distribution of the test population, cross-correlated this distribution with every template, and selected the template that yielded the highest peak normalized cross-correlation as the decoded frequency for the test sample. To ensure proper cross-validation, all test responses were removed from the template before cross-correlation. We iterated this analysis 200 times for each set of conditions. A similar procedure was followed for the array recordings, except the test set consisted of ISIs recorded in a single trial from the most highly informative channels, and the ISI templates for each frequency were constructed with all other trials of the same experimental block. We also implemented other decoders, based on Fourier decomposition or vector strength, and found these to yield comparable performance.

##### Neurometric analyses

Neurometric functions were derived in the same manner as were their psychometric counterparts. For the discrimination task, we computed the proportion of times the decoded frequency of each comparison stimulus was greater than the decoded frequency of the standard stimulus, with frequency decoded from single-trial responses. For the classification task, we computed the proportion times the frequency decoded from single trial responses was higher than the separatrix in a given experimental session. Once neurometric functions were generated, the amplitude bias was computed as were their psychophysical counterparts.

In addition, we simulated the discrimination and classification tasks with a “greater-rate” decoder, which was a proxy for basing frequency judgments solely on perceived magnitude. For the discrimination task, we computed the proportion of times the comparison stimulus evoked the higher firing rate (in single trials), averaged across the most highly informative channels; in the classification task, we computed the number of times the comparison stimulus evoked a firing rate that was higher than firing rate at the separatrix frequency, estimated by interpolating between the rates evoked by the two adjacent frequencies at the median amplitude.

##### Rate-biased timing model

We hypothesized that animals based their frequency judgments on temporal spiking patterns but exhibited some bias to select the stimulus that evoked the higher rate. To test this, we constructed a simple model where the proportion of times the animal selected the comparison as higher in frequency was a weighted sum of the prediction from the timing-based decoder and the “greater-rate” (magnitude-based) decoder:

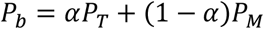

where *P*_*b*_, *P*_*T*_, *P*_*M*_, are the behavioral performance, timing-decoder performance, and magnitude-decoder performance, respectively, and α is a free parameter. Fitted α was 0.59, 0.78, and 1.02 for M4 with the 20-Hz, 50-Hz, and 100-Hz standards, respectively (the latter being the one case with a slight reversal in bias), and 0.64 for M3 with the 50-Hz separatrix.

## Acknowledgments

We would like to thank Hannes Saal and Jeff Yau for their helpful comments. This work was supported by NINDS grants NS122333, NS101325, and NS095251.

## Supplemental results

### Firing rates do not covary consistently with behavioral judgments

One way to establish a link between neural responses and behavior is to show that trial-to-trial fluctuations in the responses covary with the behavior ^45^. With this in mind, we split trials according to the binary judgments of the animals for each stimulus condition and compared the firing rates in each case (rates were z-scored within each condition be comparable across conditions (cf. ^13^). If firing rates were systematically related to behavior, we expected the distribution of firing rates to differ when conditioned on behavioral choice.

We found that population firing rate covaried significantly with behavior for both animals, but in incongruous ways. For the frequency discrimination task (monkey M3), the difference in the firing rates evoked by the two stimuli in a pair was slightly larger when the animal correctly selected the higher frequency (Figure S 15B), consistent with previous results (cf.^13^). In other words, the animal was more likely to correctly select the higher stimulus when it produced a higher firing rate. For the frequency classification task (monkey M4), however, responses to the same stimulus were slightly weaker on trials when the animal selected the stimulus as higher in frequency (Figure S 15A). This effect was observed in every frequency range, including the flutter range, over which (1) our decoding results indicate that higher rates should correspond to higher frequencies (Figure 4E) and (2) the animal demonstrated a clear bias toward higher amplitudes (Figure 1F, Figure S 1B). Therefore, while firing rates covaried with behavior in both monkeys, the relationship was inconsistent across animals, which calls into question its interpretability in the present context.

### Frequency codes are similar across somatosensory cortical fields

S1 comprises three hierarchically organized cortical fields with neurons that respond to touch, from Brodmann’s area 3b (primary somatosensory cortex proper ^46^) to area 1 then area 2 ^47^. Having previously found that progressively fewer neurons exhibit phase locking at higher stages of cortical processing ^11^, we repeated the decoding analysis with neurons split by cortical field. We found that results were generally consistent across somatosensory fields, with timing-based decoders outperforming and exhibiting less amplitude bias than rate-based ones (Figure S 13, Figure S 14). The exception was area 2, where a slight reversal in the overall performance level of the two decoders was observed, though the bias remained consistent with that observed for areas 3b and 1. One possible interpretation is that the timing signal has been converted to a rate-based on in this area, but the sample size for area 2 is too small to draw any definitive conclusions.

## Supplemental tables and figures

**Table S 1:** Summary of data sets used in this study. We collected data from 6 animals, including behavioral data (frequency classification or discrimination), neural recordings during task performance (chronic array multi-unit recordings), and neural recordings while stimuli were delivered to the animal without a behavioral task (electrode drive single-unit and array multi-unit), across many different frequency ranges. The frequency ranges of the vibrations and the figures that show the resulting analyses are listed in columns 4 and 5, respectively.

| Animals | Task | Neural recordings | Frequency range(s) | Relevant figure(s) |
| --- | --- | --- | --- | --- |
| M1 | frequency discrimination | none | 100Hz standard (50-300Hz)<br>200Hz standard (50-400Hz) | Figure 1<br>Figure S 1 A, C, D |
| M2 | frequency discrimination | none | 50Hz standard (35-65Hz)<br>100Hz standard (50-300Hz)<br>200Hz standard (50-400Hz) | Figure 1<br>Figure S 1 A, C, D |
| M3 | passive | chronic array multi-unit | 10-20Hz<br>35-50Hz<br>50-65Hz<br>100-200Hz | Figure 2A, C<br>Figure 4A-C<br>Figure S 2A-F<br>Figure S 3<br>Figure S 4<br>Figure S 5 B<br>Figure S 6 A<br>Figure S 9 |
| M3 | frequency classification | chronic array multi-unit | 20Hz separatrix (14-26Hz)<br>50Hz separatrix (35-65Hz)<br>100Hz separatrix (70-130Hz) | Figure 1<br>Figure 5<br>Figure S 1B-D<br>Figure S 5 C<br>Figure S 6 B<br>Figure S 10 A-F<br>Figure S 11 A, C<br>Figure S 12<br>Figure S 15 A |
| M4 | frequency discrimination | chronic array multi-unit | 50Hz standard (35-65Hz) | Figure 1<br>Figure 5<br>Figure S 1 A, C, D<br>Figure S 5 C<br>Figure S 6 B<br>Figure S 10 G-H<br>Figure S 11 B, D<br>Figure S 12<br>Figure S 15 B |
| M5-6 | passive | Electrode drive single-unit | 50-400Hz | Figure 2 B, D, E<br>Figure 3<br>Figure S 3<br>Figure S 7<br>Figure S 8<br>Figure S 13<br>Figure S 14 |

**Figure S 1:**
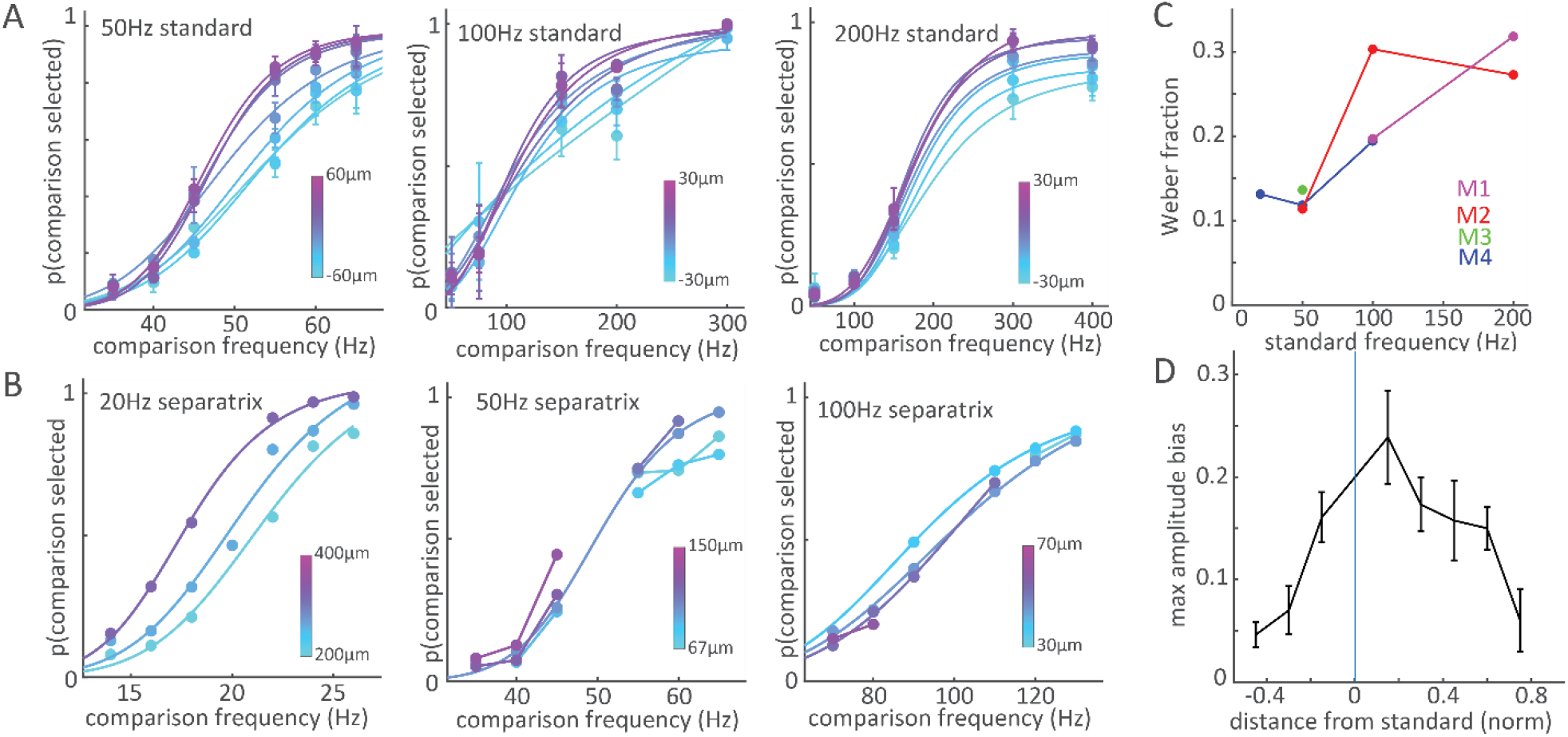
Psychophysical performance at all frequencies. A| Frequency discrimination performance for each standard frequency. Left: performance of M2 and M3 with a 50-Hz standard. Middle: performance of M1 and M2 with a 100-Hz standard. Right: performance of M1 and M2 with a 200-Hz standard. B| Frequency classification performance of M4 with separatrices of 20 Hz (left), 50 Hz (middle), and 100 Hz (right). C| Weber fractions for each animal at each standard/separatrix. D | Largest amplitude bias (spread between curves in different amplitude conditions, see Methods) as a function of the difference between the comparison and standard frequencies. The difference is normalized by the range of comparison frequencies, so data from all standards and monkeys can be pooled. Error bars denote the standard error of the mean. The amplitude bias tends to be higher for difficult comparison frequencies (those closer to the standard).

**Figure S 2.**
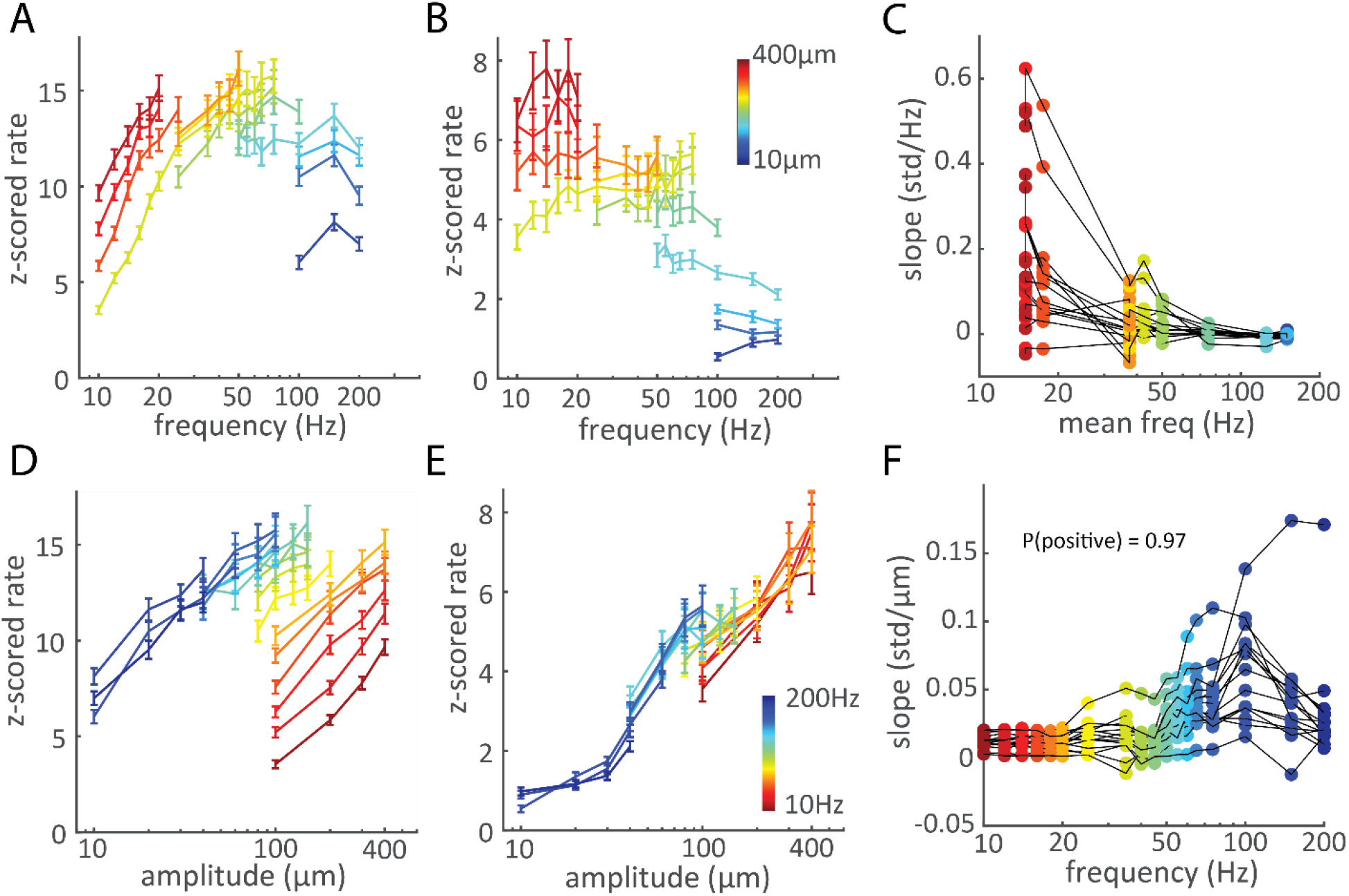
Frequency and amplitude dependence of S1 responses. A, B| Standardized firing rate versus frequency for two multi-units exhibiting different frequency-rate relationships. In one unit (A), rate increases with frequency in the low range but the slopes decrease at higher frequencies. In the other (B), the rate-frequency relationship is weak, with slopes close to zero and even negative. C| Slopes relating firing rate to frequency vs the middle frequency for the 15 most highly responsive electrodes (see Figure S 4B for responses). Each colored point corresponds to the slope of the line of matching color in panels A and B for one multi-unit. The X-coordinate corresponds to the mean frequency spanned by each of these lines. D, E| Spike rate versus amplitude for the same two electrodes in A and B. Rate rises monotonically with amplitude for both. F| Slope of the relationship between rate and amplitude as a function of frequency for the 15 most highly responsive electrodes. Each colored point corresponds to the slope of the matching line in the examples in D and E. X-axis points correspond to the frequency represented by each of these lines. The slopes across electrodes and frequencies are almost universally positive (97%).

**Figure S 3.**
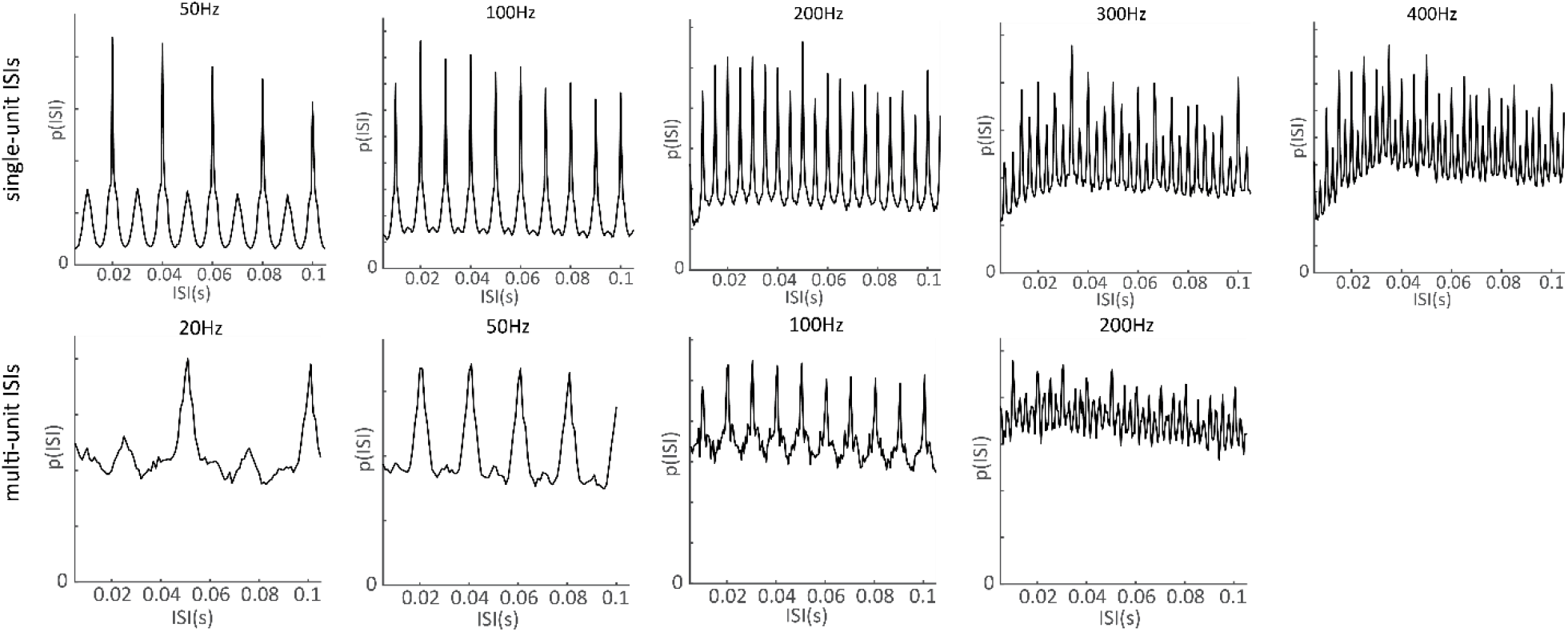
Probability distributions of ISIs in the neural population response to the indicated frequencies. The top row shows the distributions from the single-unit recordings, while the bottom row shows distributions from the array multi-unit recordings. The differences between the ISI peaks clearly correspond to the period of the stimulus frequencies. The stimulus timing is more strongly reflected in the ISI distributions of the single unit recordings as evidenced by the height of the peaks relative to the troughs. The strength of the timing declines at higher frequencies, especially for the multi-unit array recordings.

**Figure S 4.**
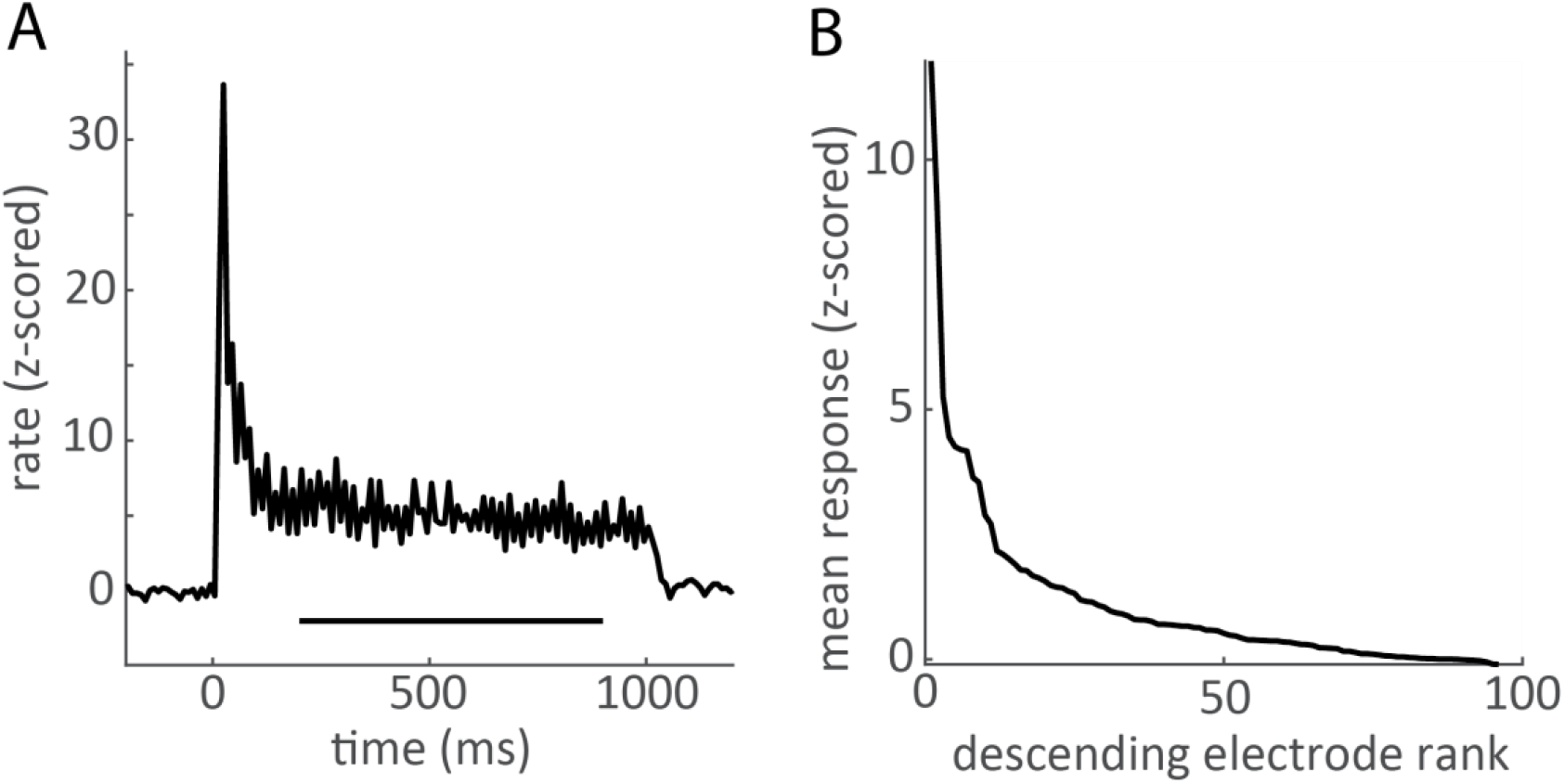
A| Peristimulus time histogram of a multiunit response to a 50-Hz vibration. Note the large transient at the onset of the stimulus (t = 0ms), a consistent feature of S1 responses to touch. For most of the analyses, we limited our analyses of rates and timing to the time interval indicated by the horizontal line. B| Responses from each M3 electrode during passive recordings sorted by overall firing rate (averaged across all stimuli). All analyses of rate were applied to the most responsive units.

**Figure S 5.**
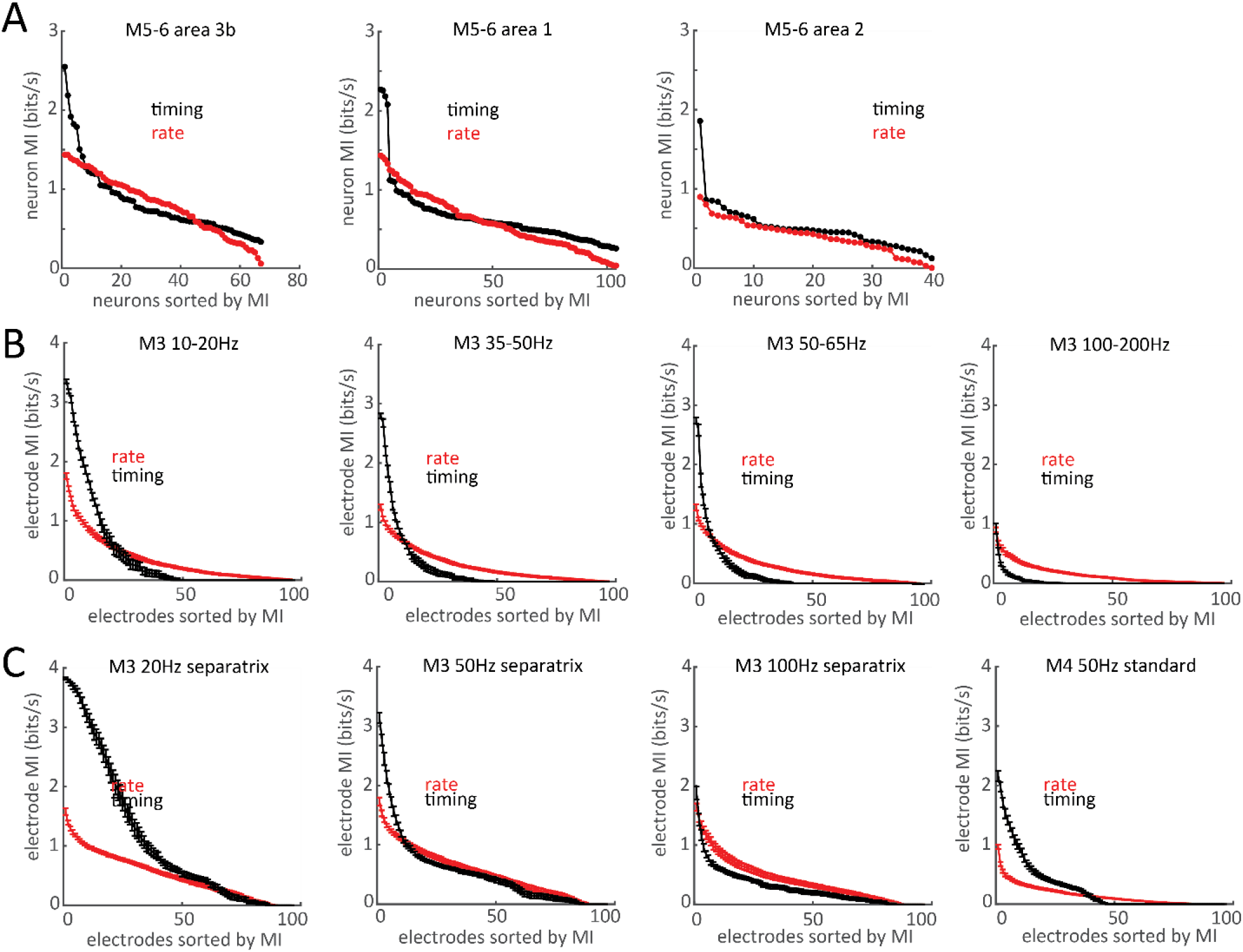
Informativeness of spike timing and rate across the neural populations. A| Mutual information between the frequency of peak-vector-strength (a measure of phase locking) and the stimulus frequency (in black), and mutual information between rate and stimulus frequency (red). The neurons are separated by cortical area and ranked according to decreasing mutual information. Note that because the peak-vector-strength frequency for a phase-locked neuron can often be a harmonic of the stimulus frequency instead of the stimulus frequency itself, this measure likely understates the information content in the timing, but functions well to find neurons with informative timing. B| For M3’s passive data in 4 frequency ranges, array channels are ranked according to mutual information between stimulus frequency and channel timing or rate. The channels are ranked with each experimental session, and error bars show the standard error across sessions. C| The same plots as in B for array recordings obtained while M3 and M4 performed their behavioral tasks.

**Figure S 6.**
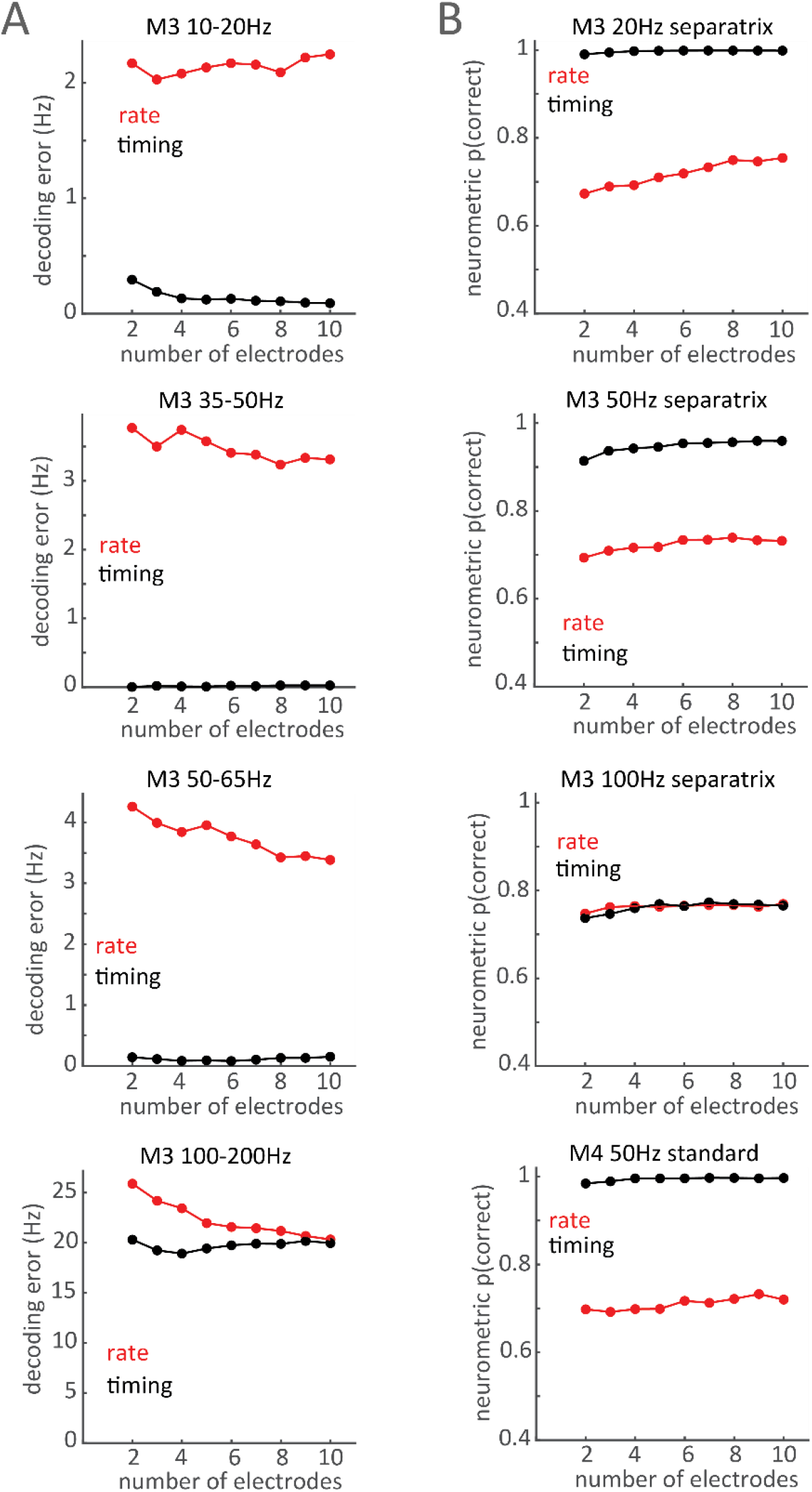
Decoder performance as a function of sample size. A | For M3’s passive recordings, decoding error as a function of the number of channels contributing to the rate decoder input (red) and the timing decoder (black). B | Neurometric performance of the rate and timing decoders (applied to the neural recordings obtained while M3 and M4 performed their tasks) as a function of the number of channels used.

**Figure S 7.**
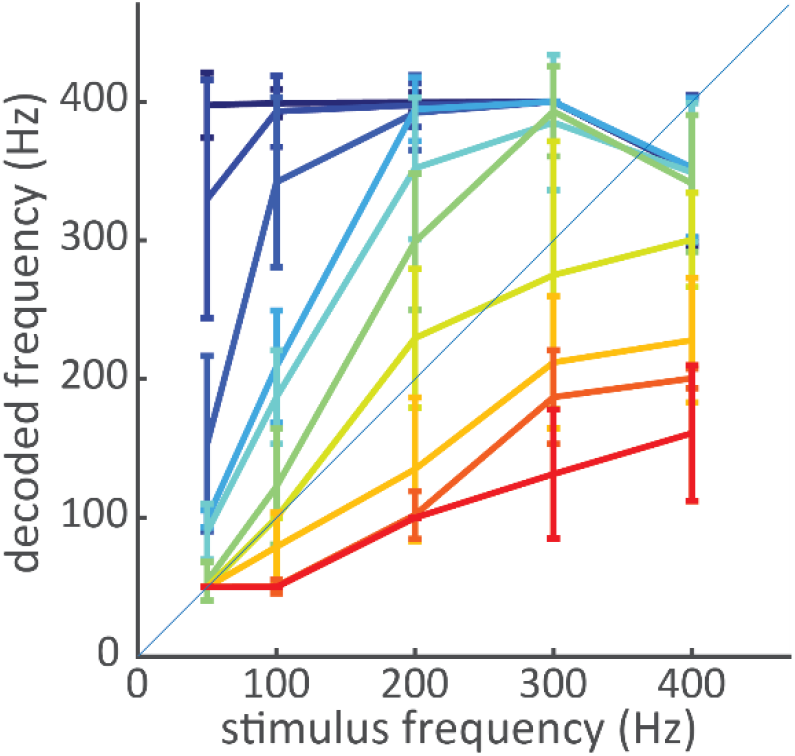
Decoding frequency from the first 250ms of the response, including the burst of activity at stimulus onset (Figure S 4A). Rate decoding is strongly biased by amplitude, as is the case when frequency is decoded from the steady-state response (200-900ms after stimulus onset).

**Figure S 8.**
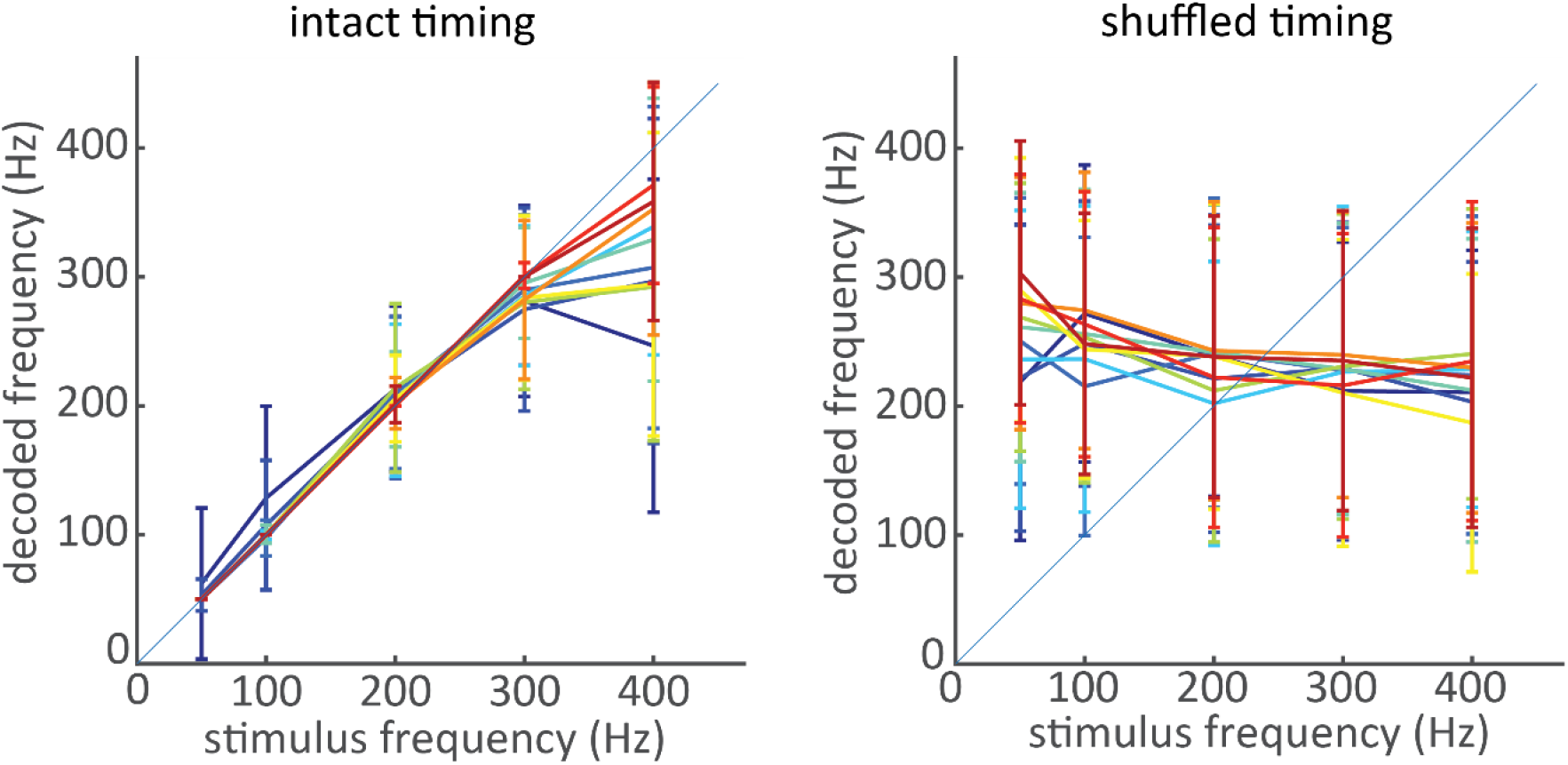
ISI-based decoders rely on spike timing. Left: when ISIs are computed from the recorded spike times, the timing decoder achieves high accuracy (same panel as Figure 3A bottom right). Right: When the spike times are shuffled before computing the ISIs, the same decoder applied to the same neural population achieves very poor performance, confirming the reliance of the timing decoder on the precise timing of spikes. The neural population here consisted of 100 randomly drawn neurons, and the error bars show the standard deviation of decoded frequency across 200 iterations.

**Figure S 9.**
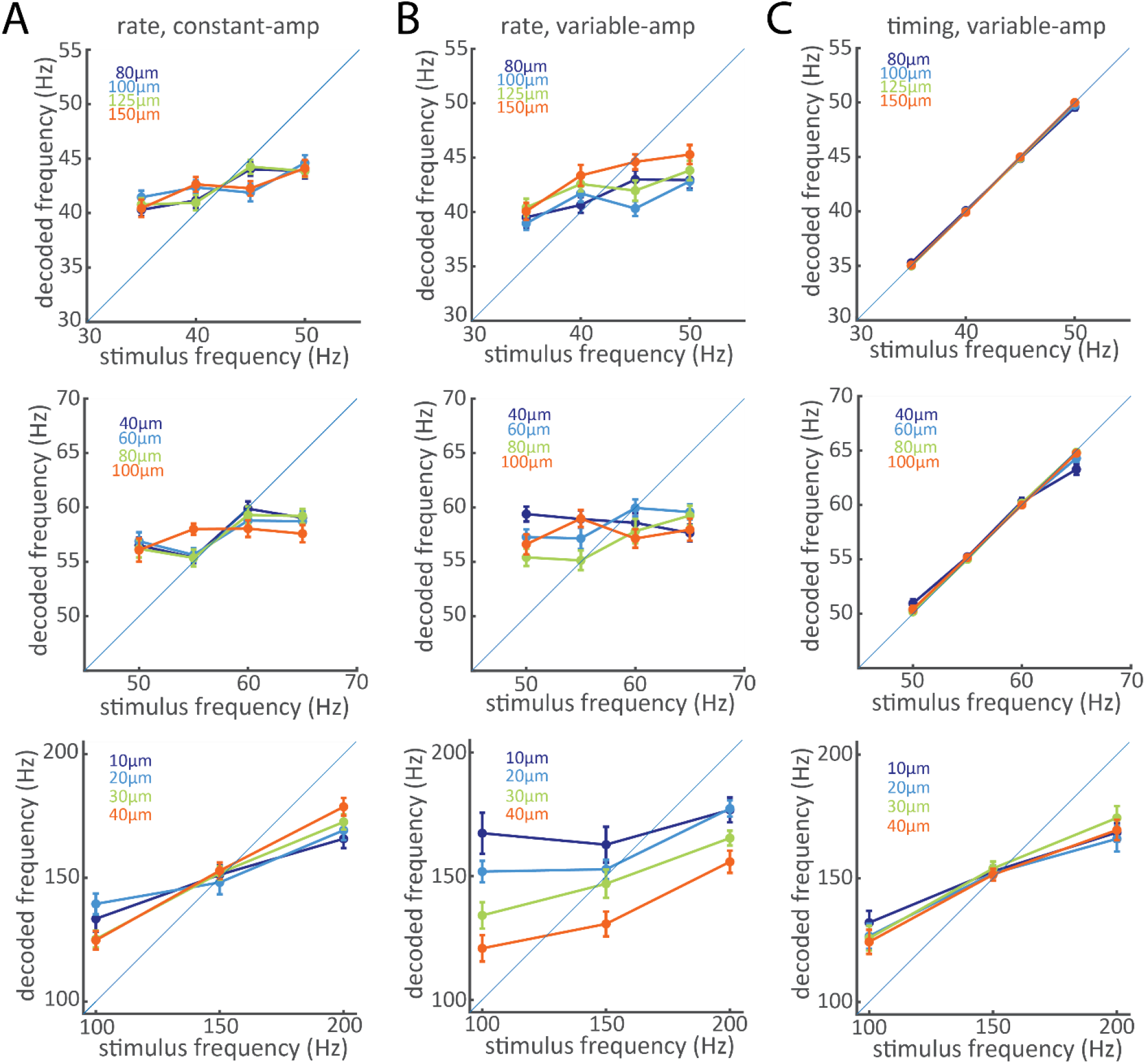
Effect of multiple amps on decoding from 30 to 200Hz from multiunit recordings from M4. Rows correspond to different frequency ranges within which the same amplitudes were used. A| Rate decoder with constant amplitudes: neural responses were grouped by amplitude and a rate decoder was trained and tested for each. B| Rate decoder with all amplitudes: a single rate decoder was trained and tested on neural responses from all amplitudes. C| Timing decoder with all amplitudes.

**Figure S 10.**
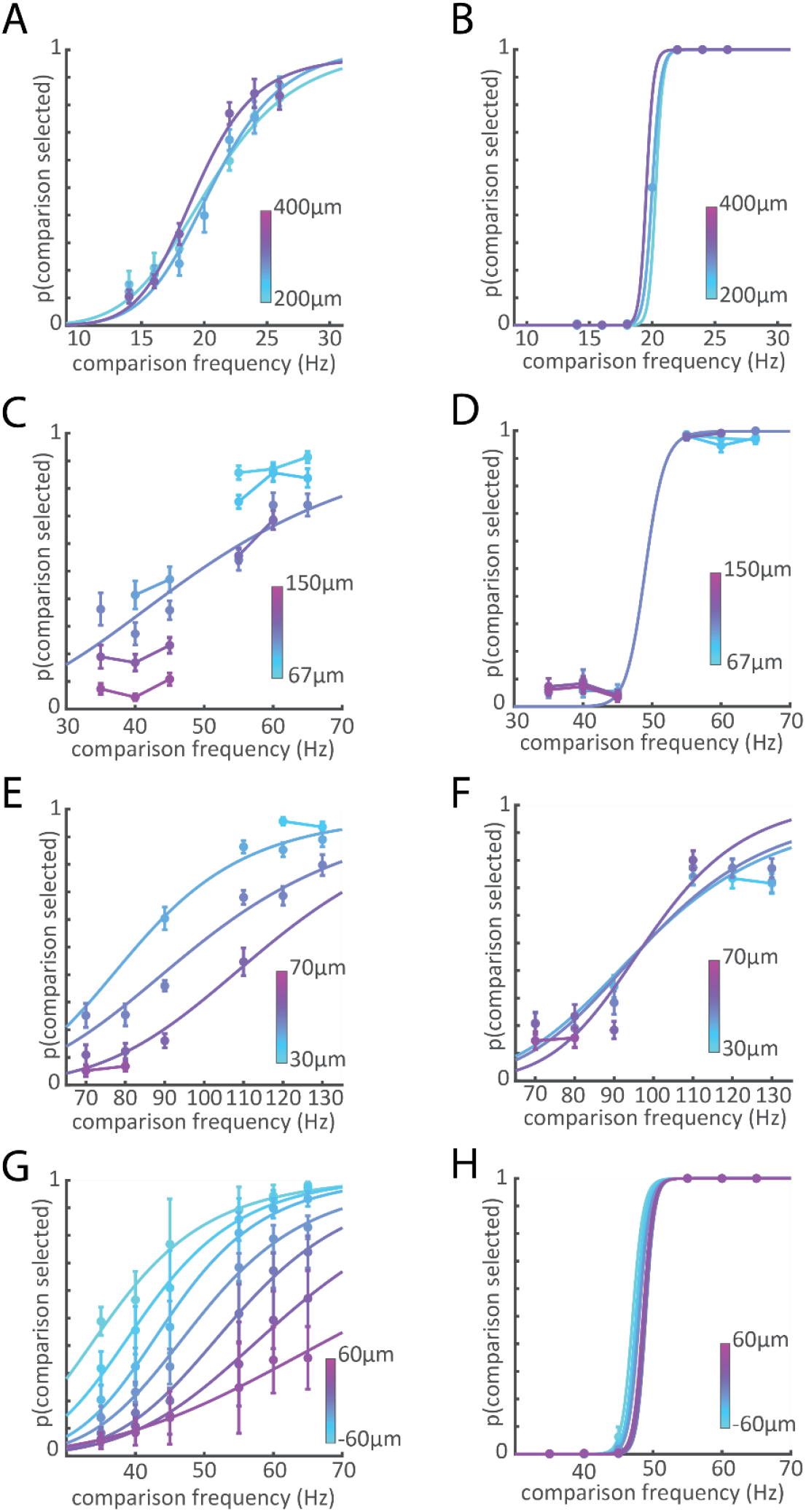
Simulation of the behavioral tasks with rate (left) and timing (right) decoders for all monkeys and standards/separatrices. A-B | Decoders applied to S1 responses from M4 as it performed the classification task with the 20-Hz separatrix. C-D| Decoders applied to responses from M4 during the classification task with the 50-Hz separatrix. E-F| Decoders applied to M4 responses during the classification task with the 100-z separatrix. G-H| Decoders applied M3 responses during the discrimination task with a 50-Hz standard (also shown in Figure 5A-B).

**Figure S 11:**
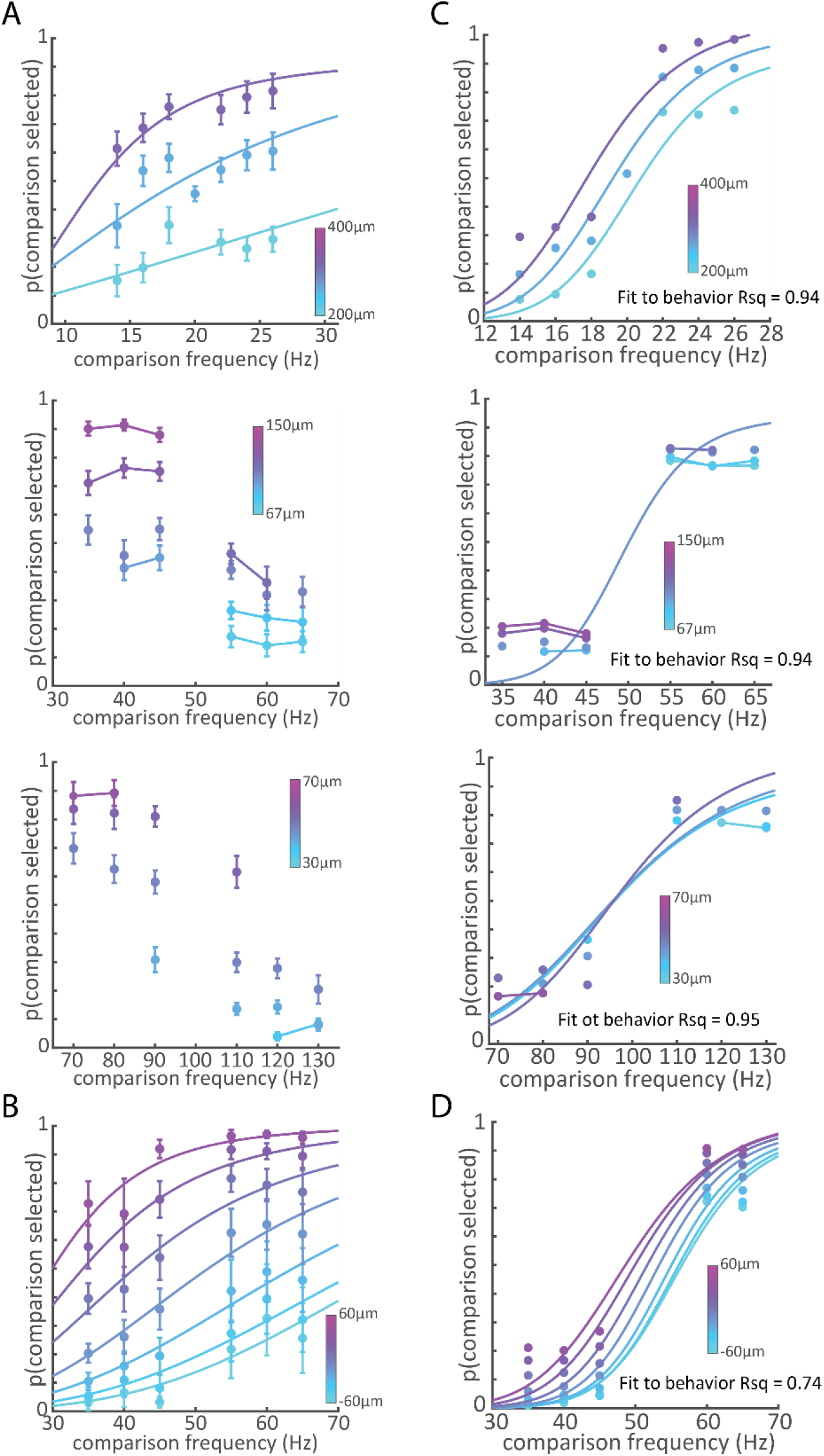
Performance of the greater-rate (magnitude based) decoder. The stimulus evoking the higher rate (averaged across inputs) is selecting as having the higher frequency. A | Neurometric performance of the greater-rate approach for M4’s responses during classification with the 3 separatrices. B | Performance of the greater-rate decoder from M3’s responses. C-D | Results of a linear model fitting the greater-rate neurometric performance and the timing neurometric performance (Figure 5B, Figure S 10 right column) to behavioral performance.

**Figure S 12.**
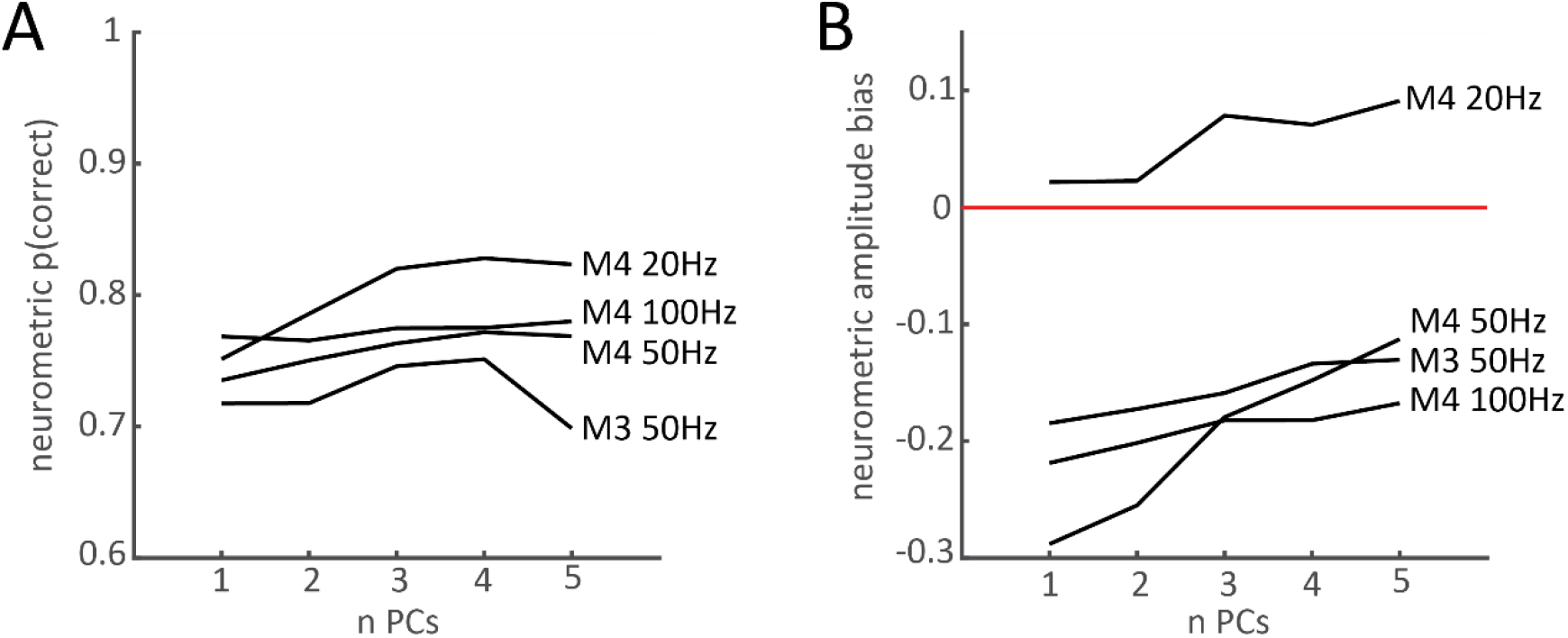
Performance of rate decoders as a function of the dimensionality of the neuronal subspace. A| Performance of the rate-based decoders as a function of the number of PCs used as decoder inputs, for each behavioral data set. B| Amplitude bias of the rate-based decoders as a function of the number of PCs. Performance and bias persist even with the addition of PCs.

**Figure S 13.**
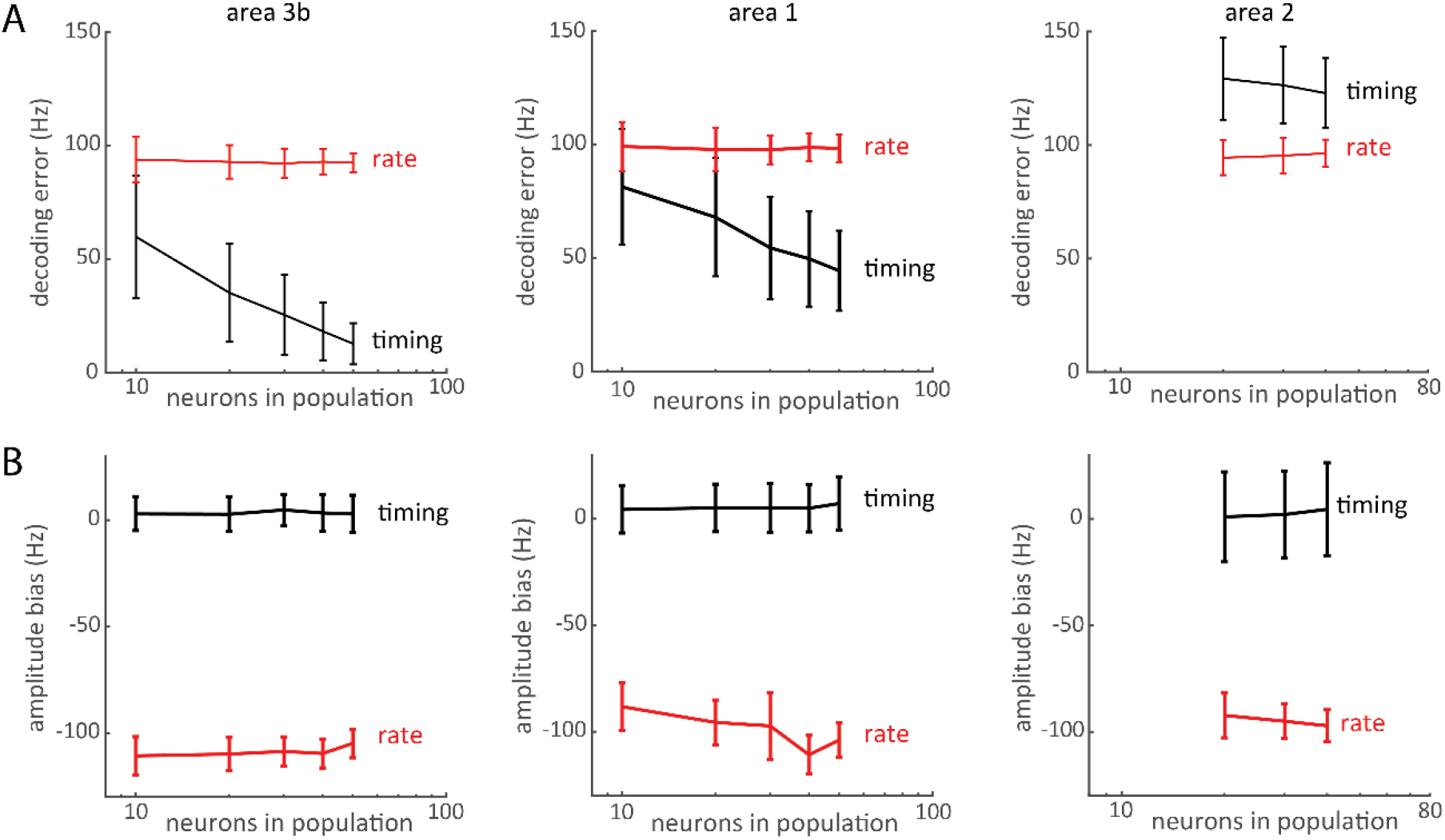
Decoding performance of rate-based and timing-based decoders with neural populations split by somatosensory cortical field. The same patterns of results were observed in all cases, except that area 2 responses yielded poorer timing-based than rate-based decoding performance, possibly reflecting a timing to rate conversion.

**Figure S 14.**
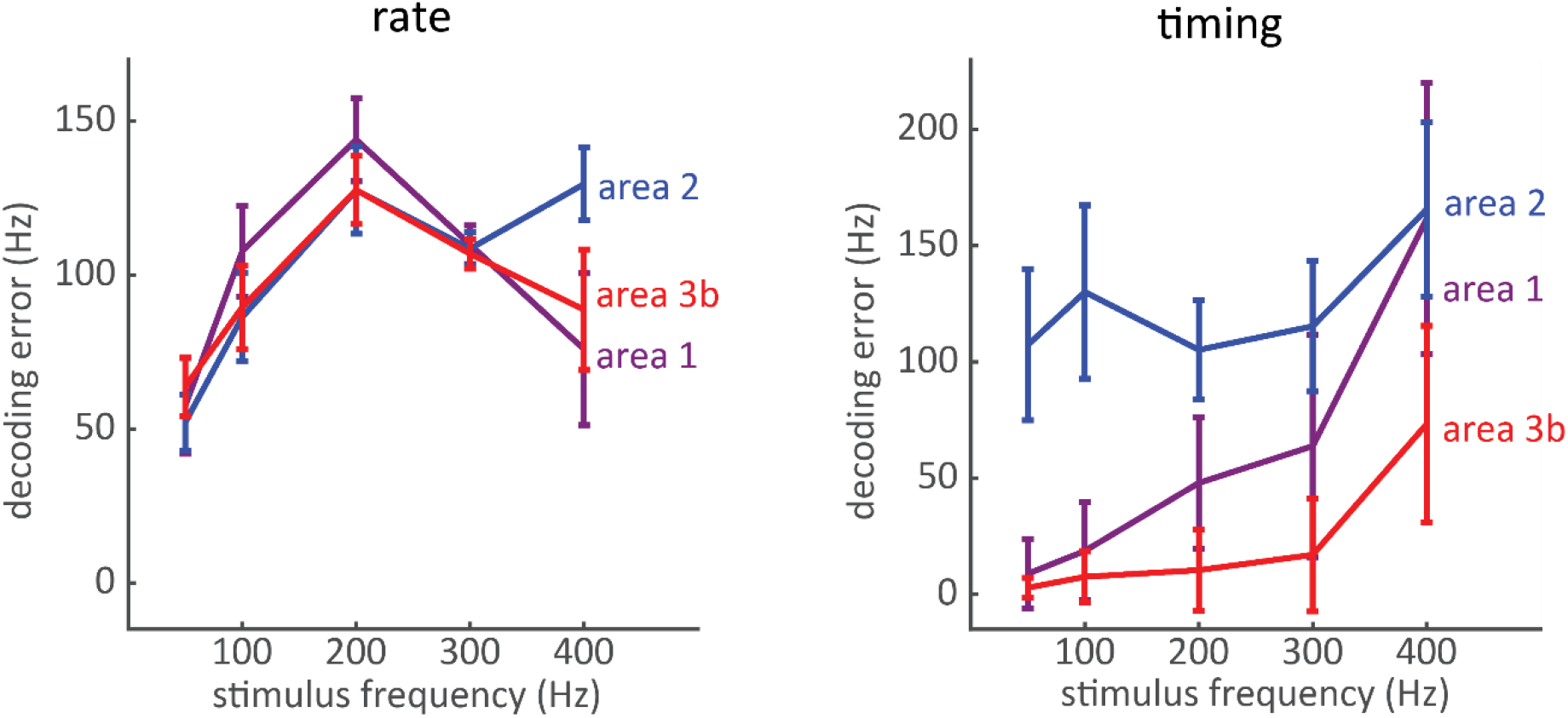
Decoding error as a function of frequency. Rate-based decoders perform most poorly around 250 Hz (the peak of tactile sensitivity) whereas timing decoders deteriorate at higher frequencies. Population responses were constructed from 40 neurons for all areas, limited by the sample size of area 2.

**Figure S 15.**
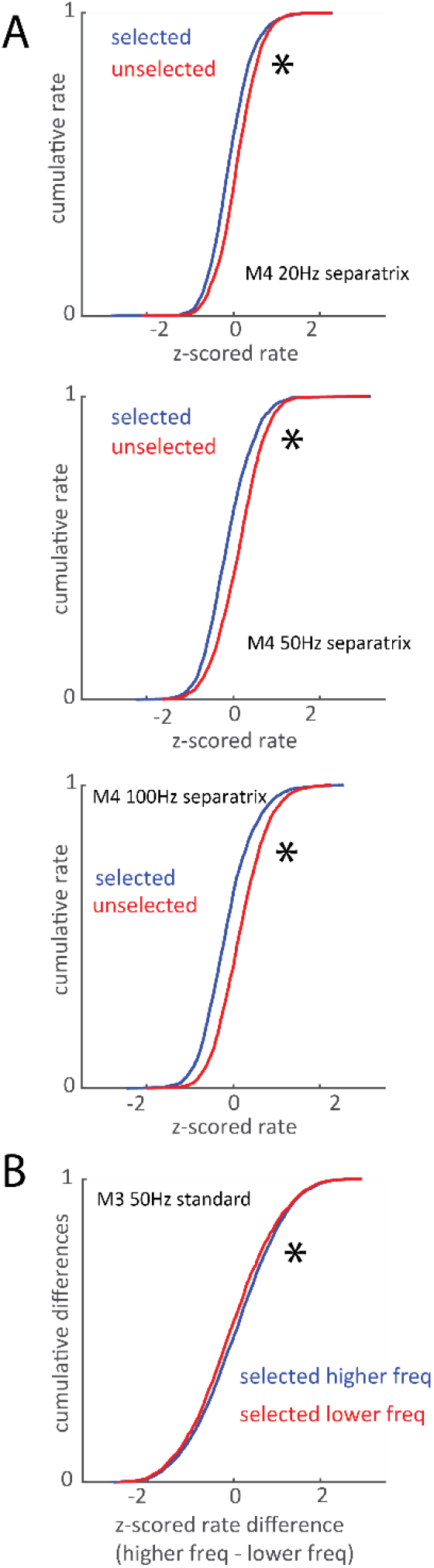
Covariance of rate and behavior, pooled across stimulus conditions. A| Cumulative distribution of firing rates across all selected (red) and unselected (blue) stimuli for M4 classification at the three separatrices. Rates were z-scored within presentations of each unique stimulus in each experimental session before being pooled. Asterisks denote significant rate differences between the selected and unselected populations (p < 0.05), with right-shifted curves corresponding to higher firing rates in the population. B| Cumulative distribution of rate differences (higher freq – lower freq) when M3 performed frequency discrimination. The difference between the two stimuli was slightly but significantly (p < 0.05) greater when the animal correctly selected the higher frequency. In other words, to the extent that the response rate to the higher frequency was greater than that to the lower frequency, the animal tended to select the higher frequency.

